# Time-resolved cryo-EM of G protein activation by a GPCR

**DOI:** 10.1101/2023.03.20.533387

**Authors:** Makaía M. Papasergi-Scott, Guillermo Pérez-Hernández, Hossein Batebi, Yang Gao, Gözde Eskici, Alpay B. Seven, Ouliana Panova, Daniel Hilger, Marina Casiraghi, Feng He, Luis Maul, Peter Gmeiner, Brian K. Kobilka, Peter W. Hildebrand, Georgios Skiniotis

## Abstract

G protein-coupled receptors (GPCRs) activate heterotrimeric G proteins by stimulating the exchange of guanine nucleotide in the Gα subunit. To visualize this mechanism, we developed a time-resolved cryo-EM approach that examines the progression of ensembles of pre-steady-state intermediates of a GPCR-G protein complex. Using variability analysis to monitor the transitions of the stimulatory Gs protein in complex with the β_2_-adrenergic receptor (β_2_AR) at short sequential time points after GTP addition, we identified the conformational trajectory underlying G protein activation and functional dissociation from the receptor. Twenty transition structures generated from sequential overlapping particle subsets along this trajectory, compared to control structures, provide a high-resolution description of the order of events driving G protein activation upon GTP binding. Structural changes propagate from the nucleotide-binding pocket and extend through the GTPase domain, enacting alterations to Gα Switch regions and the α5 helix that weaken the G protein-receptor interface. Molecular dynamics (MD) simulations with late structures in the cryo-EM trajectory support that enhanced ordering of GTP upon closure of the alpha-helical domain (AHD) against the nucleotide-bound Ras-homology domain (RHD) correlates with irreversible α5 helix destabilization and eventual dissociation of the G protein from the GPCR. These findings also highlight the potential of time-resolved cryo-EM as a tool for mechanistic dissection of GPCR signaling events.

## Introduction

G protein-coupled receptors relay extracellular signals primarily via the activation of distinct subtypes of heterotrimeric G proteins (comprised of Gα, Gβ, and Gγ subunits) that, in turn, initiate signaling cascades by interacting with downstream effectors. For the vast majority of GPCRs, agonist binding to the receptor extracellular pocket promotes conformational changes on the intracellular side, enabling the engagement of the GDP-bound Gα subunit in a G protein heterotrimer (Fig. 1a). A key player in this receptor-G protein interaction is the Gα C-terminal α5 helix, which must undergo a conformational transition to engage the receptor^1^. The repositioning of the α5 helix, in conjunction with the disengagement of the AHD from the RHD^2,3^, leads to a weaker affinity for and release of GDP^3,4^. The nucleotide-free G protein is subsequently loaded with GTP, promoting structural changes that activate Gα, weaken its affinity for the receptor, and drive the functional dissociation of the G protein heterotrimer^5-8^.

**Figure 1.**
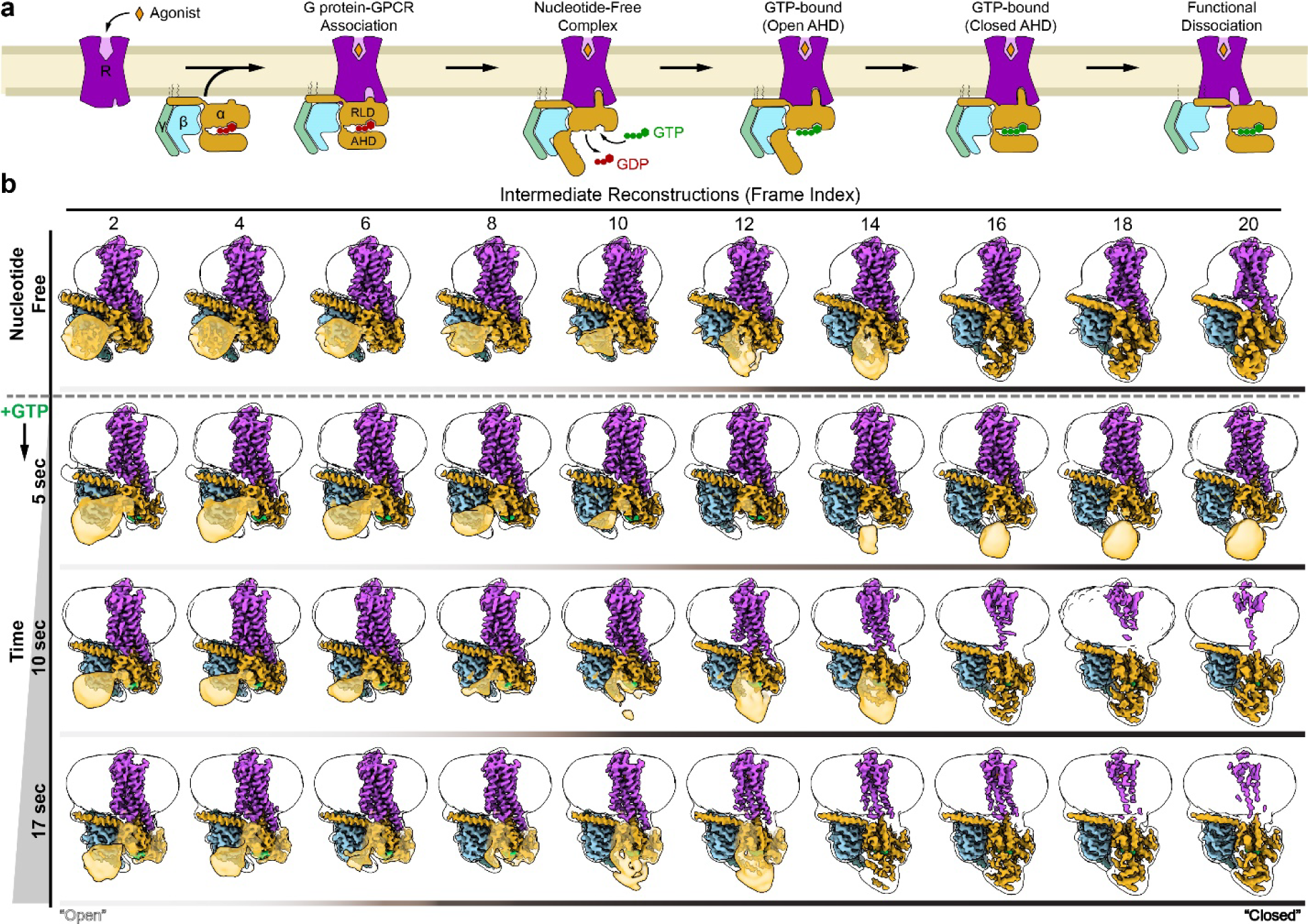
Conformational dynamics during G protein activation. **a**, In response to agonist binding, a GPCR engages heterotrimeric G protein through the Gα C-terminal α5 helix resulting in the displacement of the Gα alpha-helical domain (AHD) in relation to the Ras-homology domain (RHD). This opening allows for the release of bound GDP and the subsequent binding of GTP, leading to Gα subunit activation and functional dissociation of Gβγ from Gα. **b**, β_2_ΑR-Gs conformational dynamics revealed through cryo-EM. Complexes were captured by vitrification in the nucleotide-free state. Utilizing cryoSPARC’s 3DVA function, the data was divided to obtain 20 reconstructions across the major principal component of motion (*i.e.*, AHD closure). For space consideration, only even numbered intermediates (frame indexes) are shown. Complexes were additionally frozen at progressive time-points following the addition of GTP to the nucleotide-free complex (last 3 rows). Using the same processing schema, the dynamics of the GTP-bound complex revealed the proportion of particles with a closed AHD to increase with increasing times of vitrification post-addition of GTP. Reconstructions shown include the sharpened maps in solid coloring, surrounded by the gaussian filtered unsharpened envelope to show the micelle and location of the AHD (translucent gold, except when observed directly in sharpened map). Color bars beneath each structural ensemble are shaded in relation to the observation of ‘open’ or ‘closed’ AHD position.

Although the pathway from receptor agonism to G protein activation is a dynamic, multi-step mechanistic process^3,9-13^, structural studies have been limited in their inability to capture different sub-states. Since the initial crystal structure of β_2_AR in complex with Gs protein^1^, the advent of cryo-electron microscopy (cryo-EM) has facilitated countless structures of GPCR-G protein complexes^14-16^, providing a wealth of information on ligand recognition, receptor activation, and G protein coupling under nucleotide-free conditions. The G protein has the highest affinity for the receptor in the absence of nucleotide, and thus nucleotide-free conditions have been invariably used for structural studies to promote receptor complex stability, which is often further enhanced with stabilizing nanobodies or antibodies^15,17^. However, given the constant presence of nucleotides in the cytoplasm, a nucleotide-free GPCR-G protein is likely extremely transient *in vivo*, suggesting that these structures provide a very narrow window into the G protein activation process. Yet to be captured are short-lived transition intermediates associated with G protein coupling, GDP release, and GTP binding leading to activation of the G protein heterotrimer and its functional dissociation from a GPCR. Such structural information is critical to outline the conformational landscape of the dynamic GPCR signaling systems, understand the basis for G protein selectivity^18^ and evaluate the effects of drugs with distinct efficacies and potencies to enable more rational pharmacology^19^.

To begin addressing this limitation, we sought to visualize by cryo-EM the transition of conformational ensembles of β_2_AR in complex with heterotrimeric Gs protein after adding GTP. The β_2_AR is a Family A GPCR that primarily couples to Gs to increase intracellular cAMP levels^20^, thereby regulating crucial physiological responses, such as smooth muscle relaxation and bronchodilation^21-23^. The β_2_ΑR-Gs signaling system has been historically well-studied, providing various lines of biochemical, biophysical, and structural data that can support mechanistic investigations^1,5,9-11,24-26^. Our early EM analysis of β_2_ΑR-Gs upon negative-stain “fixation-trapping” on EM grids within several seconds after the addition of GTPγS^5^, revealed distinct complex dissociation intermediates. Even though at low resolution, that work provided a valuable demonstration that such direct visualization is feasible without pursuing sample mixing and freezing at the msec scale. Inspired by these studies, here we employed cryo-EM and “freeze-trapping” at distinct time points after the addition of GTP to examine ensembles of β_2_ΑR-Gs complex and reconstruct multiple ordered states from conformationally heterogeneous complexes. By monitoring how distinct structural populations evolved over time compared with ‘checkpoint’ crystal structures, we were able to capture, in high resolution, the temporal ordering of events underlying G protein activation on the receptor. This time-resolved cryo-EM approach to visualize pre-steady state β_2_ΑR-Gs-GTP intermediates presents both opportunities and challenges for exploring key molecular recognition events underlining the highly tuned GPCR signaling mechanisms.

## Results

### Conformational dynamics of the nucleotide-free complex

In a first study, we evaluated the dynamic behavior of detergent-solubilized nucleotide-free β_2_AR-Gs complex (β_2_AR-Gs^EMPTY^) by cryo-EM, further aiming to establish a baseline for complex stability under these conditions. To capture the full dynamic range of complex conformations, we chose not to employ any scFv or nanobody stabilizers. Instead, we enhanced sample stability by activating the receptor with c-Epi, a conformationally-constrained epinephrine, that is a highly efficacious and β_2_AR-selective agonist. Prior studies, including our earlier EM work with negative-stained particles revealed the dynamic positioning of AHD in the β_2_AR-Gs^EMPTY^ complex^3,5^. Similarly, in our current cryo-EM study, a conventional three-dimensional classification approach shows different locations of the AHD (Supplemental Fig. 1a). To explore these conformations and their transitions, we employed 3D variability analysis (3DVA)^27^ as implemented in cryoSPARC, which separates and orders projections based on particle conformation along principal components of variability, thereby enabling a view of the main directions of macromolecular dynamics observed in a complex (Fig. 1, Supplemental Fig. 1). The first principal component (PC0), split amongst twenty frames with overlapping particles, shows an extensive swing-like movement of the density corresponding to the AHD between a fully open and a fully closed position against the RHD. Of note, there are predominantly two overall locations of the AHD, open versus closed, with limited occupancy of transitions between them (Fig. 1b, Supplemental Figs. 1-2). By contrast, the rest of the complex along this primary principal component appears overall conformationally stable.

To better understand the underlying structures, we extracted twenty sequential sets of overlapping particles along PC0, which were used to refine twenty transitionary cryo-EM reconstructions with global indicated resolutions between 3.2Å – 4.2Å (Fig. 1, Supplemental Fig. 1). In the most open conformations, the cryo-EM density of the AHD pivots away from the closed position by ∼61° and lies adjacent to the 2^nd^ and 3^rd^ propeller blades of the Gβ subunit (Supplemental Fig. 2). This is different to its position in the β_2_AR-Gs crystal structure (Supplemental Fig. 2h), where the AHD is further pivoted away from the RHD (∼88°) to enable its interaction with the 1^st^ and 2^nd^ blades of the β-propeller, a difference that could arise, at least in part, from crystal packing. The cryo-EM structure of NTSR1-Gi (PDB:7L0Q)^28^ also resolves the open Gαi AHD adjacent to the 2^nd^ and 3^rd^ Gβ blades, although seemingly in a distinct orientation from that of the β_2_AR-Gs cryo-EM structure, a deviation that mostly stems from differences in the Gα subtype. The analysis of the conformational variability of the β_2_AR-Gs complex in its nucleotide-free form provided a baseline to compare the conformational dynamics of the complex under all other conditions probed in this study. Nevertheless, in a cellular context, the nucleotide-free state is unlikely to exist for any significant length of time, as the high concentration of GTP (∼300 μM, compared to ∼36 μM GDP^29^) in human cells drives immediate nucleotide binding with subsequent G protein activation and functional dissociation from the receptor^5^.

### Sequential freeze-trapping for time-resolved cryo-EM of β_2_AR-Gs-GTP

To visualize the molecular changes leading to G protein activation and functional release from the receptor upon nucleotide binding, we developed a time-resolved cryo-EM approach whereby we vitrified and imaged β_2_AR-Gs complex at short sequential time points (5 sec, 10 sec, and 17 sec) post addition of GTP. The 3DVA analysis, as implemented above, revealed a range of complex conformations analogous to the nucleotide-free complex but with two notable differences: First, the population of particles with a closed AHD conformation increases progressively with the time of GTP incubation prior to freeze-trapping. Second, the later frames in the trajectories for 10 sec and 17 sec show disappearing receptor densities, suggesting complex destabilization (Fig. 1b, Supplemental Figs. 2-5).

To verify that our observations resulted from properly ordered structural transitions and to classify the conformers from different time points within the same PCA trajectory, we merged the curated β_2_AR-Gs^GTP^ particles of all time-points together, and processed this larger dataset by 3DVA to obtain twenty ordered cryo-EM reconstructions from overlapping particle distributions with global indicated resolutions of 2.9Å – 3.6Å (Fig. 2, Supplemental Fig. 6). Like the β_2_AR-Gs^EMPTY^ and individual β_2_AR-Gs^GTP^ datasets, we observed that the position of the AHD remained the most recognizable primary variable across the trajectory, proceeding from an open AHD conformer to a closed AHD conformer (Supplemental Fig. 2). Moreover, when each intermediate reconstruction frame was analyzed to determine the origin of particles, it became apparent that projections from our shortest time point (5 sec) contributed more to the frames with an open AHD (early intermediate reconstructions), with minimal contributions to late frames in the trajectory. By contrast, as the conformers progressed to a closed AHD position (ordering from intermediate 1 to 20) we observed increasing contribution from the later time-point datasets (*i.e.*, 10 sec followed by 17 sec) (Fig. 2b). The expected distribution of particles from individual datasets with increasing time across the combined trajectory supports the relative robustness of our approach despite the limited features of the relatively small membrane protein complex. Furthermore, the merging of datasets enabled us to increase the number of projections contributing to every conformation, potentially improving the projection classification and also the resolution of each intermediate map. Combined with the structural comparisons to known structures detailed below, these results further enhanced our confidence that the conformational transitions that have driven the 3DVA trajectory stem from temporal, coordinated dynamics rather than stochastic motions following the addition of GTP.

**Figure 2.**
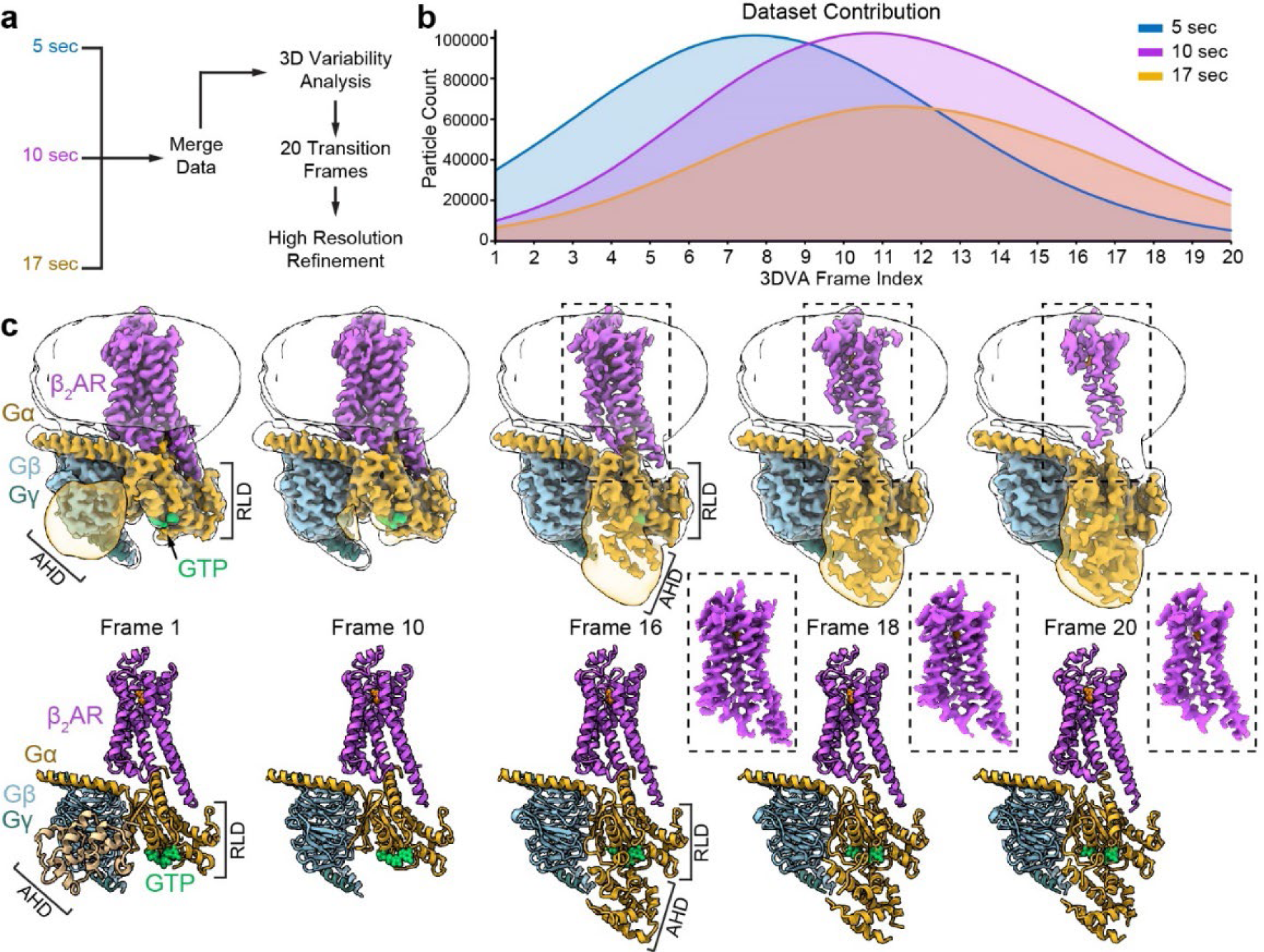
Changes in Gα structure initiated by GTP binding. **a**, Individual, curated β_2_AR-Gs^GTP^ datasets were combined and processed together to produce a consensus 3DVA trajectory. **b**, A query of the contribution of individual datasets to each intermediate reconstruction (frame) revealed that early intermediates (open AHD) are comprised primarily of particles from the earliest time point (5 sec), and later time points (10 and 17sec) correspond increasingly to later intermediate reconstructions (closed AHD). **c**, Selected cryo-EM reconstructions (top) and models (bottom) resulting from merging β_2_AR-Gs^GTP^ datasets. Note that (1) density for GTP (green) is clearly present across the entire trajectory, (2) as the AHD domain transitions to a closed conformation and becomes more stabilized (*i.e.*, density appears) the density for the transmembrane helices of β_2_AR appears progressively weaker at the same contour level, suggesting flexibility of the receptor as it relates to the G protein heterotrimer. Local refinement of the receptor density alone produces maps with stable features throughout the 7TM, shown in dashed boxes.

Consistent with diffusion limited binding of nucleotide to the G protein, density for GTP is clearly observed within the nucleotide-binding pocket across all frames in the 3DVA trajectory, but the AHD becomes stabilized into a closed conformation only in later frames (Fig. 2, Supplemental Figs. 2, 6, and 7). From a cursory vantage point, the β_2_AR-Gs^EMPTY^ and β_2_AR-Gs^GTP^ trajectories appear similar in the AHD motion from an ‘open’ to a ‘closed’ position (Fig. 1b and Supplemental Fig. 8). However, the positioning of the ‘closed’ AHD in relation to the RHD is different in the presence and absence of nucleotide, as is exemplified in the comparison of the angle of the AHD helix αA (17° deviation) (Supplemental Fig. 8). Because in the nucleotide-free state the AHD samples both the open and closed states relatively equally (Supplemental Fig. 2), we infer that the binding of GTP does not allosterically trigger AHD closure; rather, the presence of GTP locks the AHD against the RHD domain as the AHD stochastically samples the closed conformation. Reciprocally, the fully closed AHD promotes further stabilization of GTP within the nucleotide-binding site, with the nucleotide participating in salt bridge interactions between the AHD and RHD.

The AHD must be open for the initial binding of nucleotide to the RHD^3,5^, and our maps collectively suggest that GTP can remain engaged to its binding site without the immediate closure of the AHD. This observation is consistent with studies using non-functional constructs of Gα lacking the AHD but still able to bind nucleotide, albeit unproductively or with diminished activity^30-32^, as well as studies of nucleotide binding in other small GTPases lacking a helical domain (*e.g.*, Ras, Rab, Rho)^33^. Significant changes in the RHD and its interaction with the receptor occur only after the AHD has closed. One of the most striking observations of our analysis is that the ordering and full closure of the AHD correlates with a decrease in resolvable density of the β_2_AR transmembrane region (Fig. 2, Supplemental Fig. 6). Notably, this phenomenon is not observed in the structures of the nucleotide-free complex, suggesting a significant change in interactions between receptor and G protein in response to G protein activation by GTP.

### Sequential G protein rearrangements in response to GTP loading

The cryo-EM maps from overlapping particle subsets across the variability trajectory of the combined dataset enabled us to generate twenty structures representing GTP-driven transitions (Fig. 2c). To further investigate how the binding of GTP at the nucleotide-site triggers G protein activation and disengagement from the receptor, we analyzed the dynamic structural events occurring across these structures. Starting from the GTP binding site, we observe that in initial frames with a fully open AHD, the phosphate tail of GTP maintains weak interactions with residues of the α1 helix and the highly conserved P-loop^34^ (β1-α1) of the Gαs RHD, while the GTP purine ring is stabilized through backbone contacts with the TCAT loop (β6-α5) and the hinge between the β5 strand and αG helix (Fig. 3, Supplemental Fig. 7). The TCAT loop connects the β6-strand to the α5 helix, which is the primary G protein element engaging the receptor. As the transition progresses, the GTP phosphate tail becomes further stabilized by the P-loop with an associated translation of the nucleotide by ∼2Å within the binding pocket (Fig. 3), and a corresponding change on the conformation of the TCAT loop that follows the movements of the purine ring (Fig. 4). The stabilization of GTP-P-loop interactions correlates with an extension by 1.5 helical turns of the α1 helix, which directly connects to the AHD. This extension of the α1 helix seems to require the presence of nucleotide, as it is not observed in the β_2_AR-Gs^EMPTY^ trajectory. Notably, in the nucleotide-free complex the RHD elements (*e.g.*, α1, α5, TCAT loop) do not undergo any conformational changes as the AHD progresses from open to closed conformation but instead maintain the same position to the one observed in the nucleotide-free crystal structure (PDB:3SN6) (Fig. 4, Supplemental Fig. 7).

**Figure 3.**
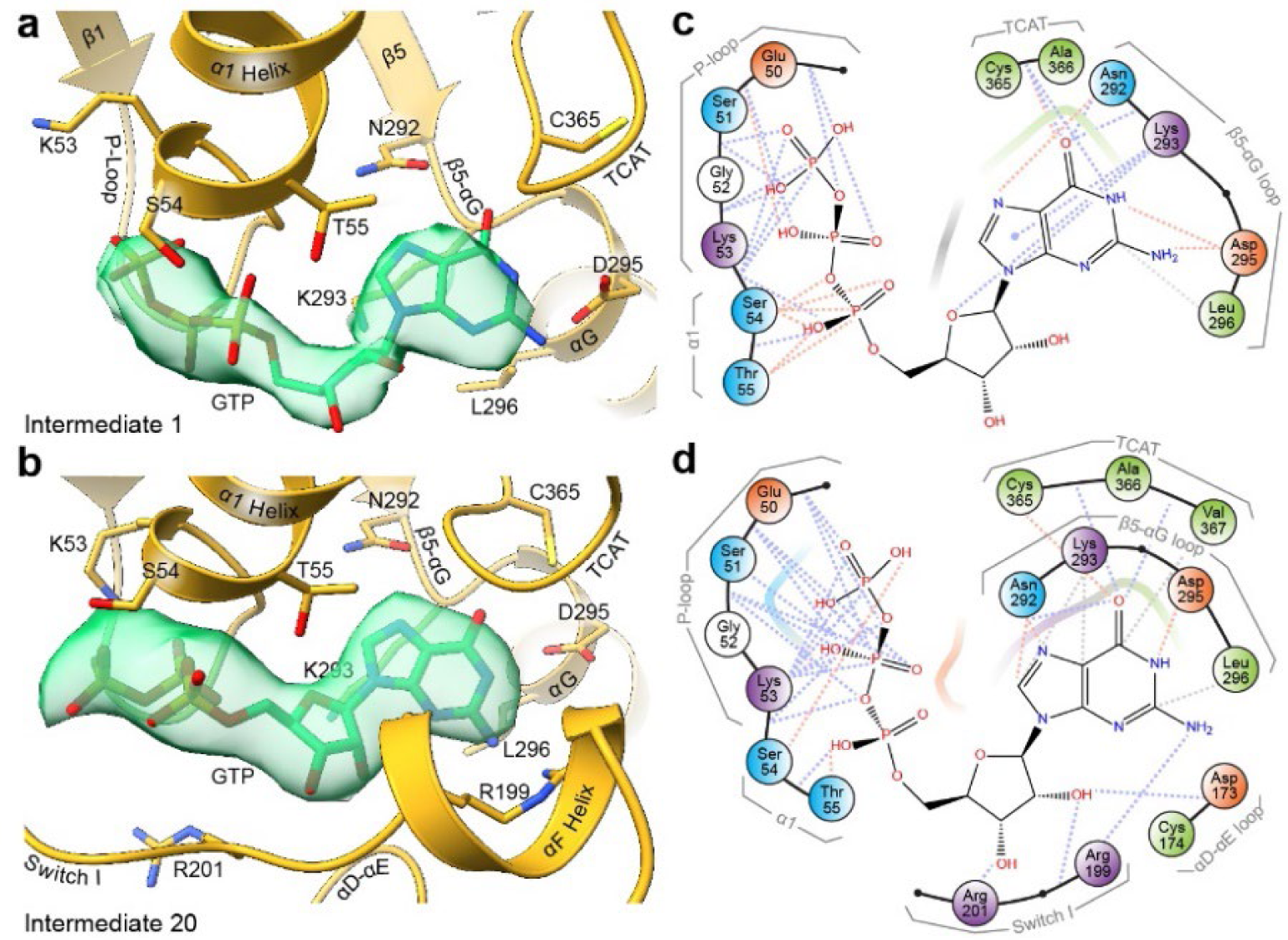
GTP transitions from loose interactions to robust stabilization within the nucleotide binding pocket. **a, b,** Comparison of the GTP binding site between the first, **a**, and last, **b**, intermediates resolved by 3DVA analysis. Cryo-EM density for modeled GTP is shown in translucent green. **c, d**, Schematic representation of GTP contacts in intermediate reconstructions 1, **c**, versus 20, **d**. Initial interactions of the GTP phosphate tail with the P-loop are strengthened over the course of the trajectory, while interactions with the TCAT and β5-αG loop become weakened and contacts form with Switch I and the αD-αE loop of the AHD.

**Figure 4.**
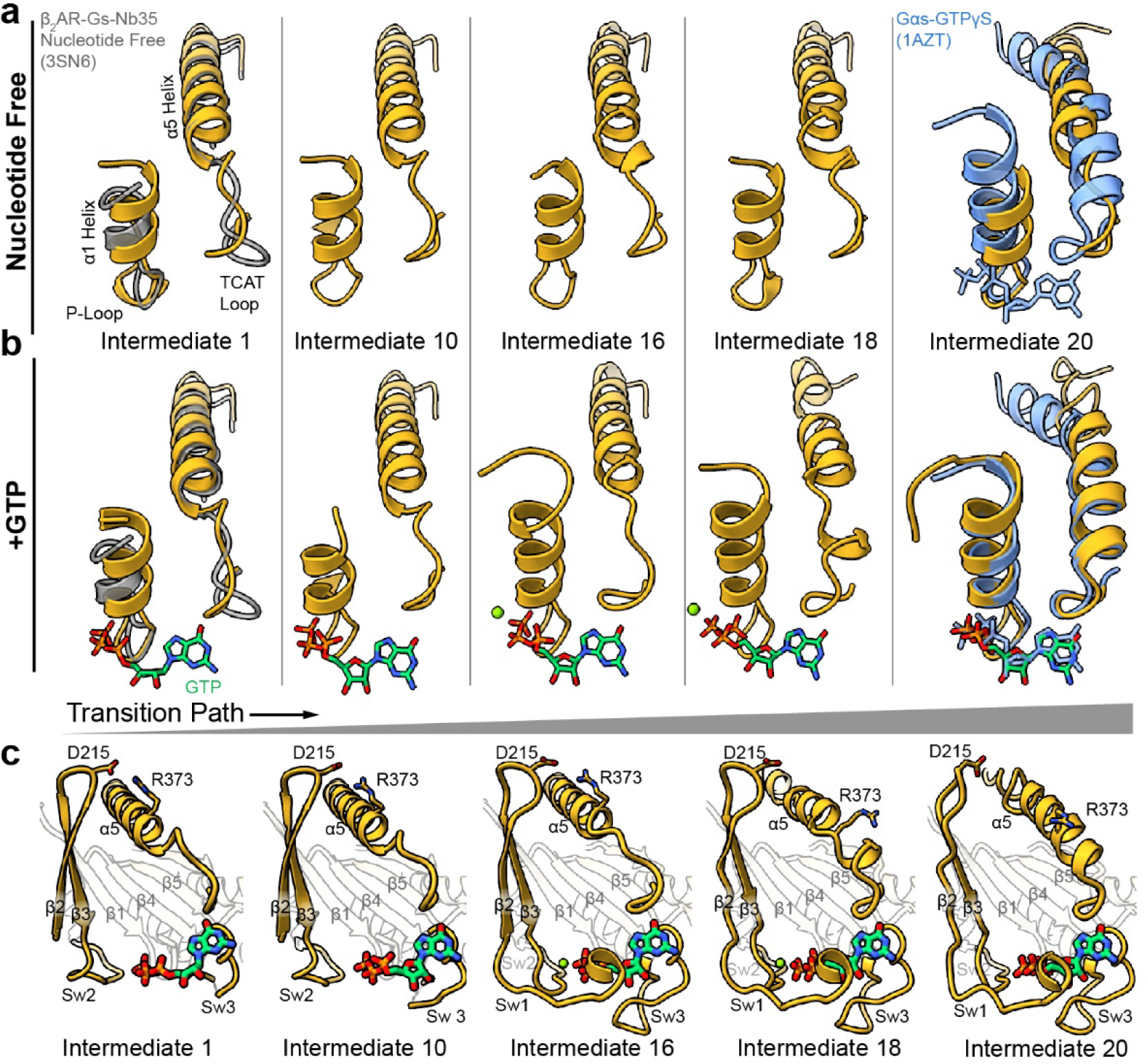
Cryo-EM structures reveal transition intermediates between steady-state structures of nucleotide free Gαs and activated Gαs-GTPγS. a, Closure of the AHD alone does not promote notable changes to the RHD elements (α1 and α5). **b,** In contrast, the presence of nucleotide induces movement of the TCAT motif and extension of the α1 helix. **c,** Over the transition path to activation, the Switch regions (I-III) become stabilized towards the nucleotide binding site. An ionic lock between the β2-β3 loop and α5 helix breaks as the α5 helix shifts to form with a new register closer to the TCAT loop.

The Switch regions (SwI-III) of the Gα RHD undergo conformational transitions during activation to facilitate GTP binding and target downstream effector enzymes, primarily adenylyl cyclase in the case of Gαs^6,35^. Following closure of the AHD, initial stabilization of the GTP phosphate tail and α1 helical extension, SwII begins changing conformation to orient towards the nucleotide binding pocket, while SwIII, which is not fully resolved in early intermediates, starts to order towards the nucleotide, likely due to contacts formed with the αD-αE loop of the closed AHD (Fig. 3). The repositioning of SwII closer to GTP correlates with the observation of a Mg^2+^ density that is partially coordinated by the phosphate tail of GTP, Asp223 of SwII, and Thr204 of SwI. The short loop connecting the Gαs β2-β3 strands, lying between the SwI and SwII regions, contains an aspartic acid residue (Asp215) that forms an ionic interaction with Arg373 on the α5 helix of Gα in the early intermediate conformers (Fig. 4c). This interaction helps stabilize the α5 helix in its extended conformation towards the receptor. In the later frames of the conformational trajectory, the movement of SwII correlates with the movement of the β2-β3 linker, in a lever-like fashion, away from the α5 helix. This separation, in conjunction with a loss of helicity in α5 near the TCAT motif, breaks the Asp215-Arg373 interaction and the helical register of α5 (Fig. 4a), and allows for the reformation of a new register where α5 begins three amino acids earlier, bringing it a helical turn closer to the TCAT motif. The change in helicity also displaces the α5 residue Phe376, previously identified as a relay during activation^36^, from interacting with β_2_AR Phe139^34.51^ (Ballesteros-Weinstein^37^ numbering in superscript) within intracellular loop 2 (ICL2), thus losing a critical interaction with the receptor. In the new α5 helical register, Phe376 is moved backward and protected by a hydrophobic groove of the RHD β-sheets (Supplemental Fig. 8). Most notably, except for the most C-terminal portion of α5 that has not fully formed into a stable helix, the RHD elements within the final intermediate structure are strikingly similar to those observed in the crystal structure of the activated Gαs-GTPγS structure (PDB:1AZT)^38^ (Fig. 4, Supplemental Fig. 8). The observation that our trajectory reveals a transition series from a conformation with open AHD where the G protein assumes a structure like the crystal structure of Gs bound to β_2_AR (PDB:3SN6)^1^ to a conformation with closed AHD in which the receptor-bound Gαs subunit is near identical to the crystal structure of the activated G protein alone (1AZT)^38^ strongly supports that, within the limitations of a linear subspace fitting of our data implemented in 3DVA, these reconstruction frames reflect an appropriately ordered chain of events leading to G protein activation after GTP binding.

### Destabilization of the GPCR-G protein interface

Also observed in the later intermediates of the cryo-EM trajectory is a decrease in observable density corresponding to the β_2_AR transmembrane helices. This may have resulted from a number of factors, such as flexibility in the association between receptor and G protein, increased plasticity in 7TM helices or even partial occupancy resulting from a fully dissociated complex. Our 2D classification analysis of the projections contributing to the final reconstruction (Intermediate 20) uniformly presented density for the receptor in detergent micelle (Supplemental Fig. 9a), suggesting that the decrease in 7TM resolvability resulted from flexibility rather than dissociation. To resolve whether the observed increase in β_2_AR flexibility arose from a rigid body motion of β_2_AR or flexibility within individual 7TM helices, particles from each intermediate reconstruction were subjected to local refinement of the receptor density, producing cryo-EM maps with indicated resolutions between 3.2Å – 4.1Å (Fig. 2, Supplemental Figs. 6 and 9). The local receptor reconstructions for frames #18-20 were highly similar at the secondary structure level, and compared to earlier frames exhibited mostly minor movements in residue side chains and a small movement of the ligand towards ECL2/TM2 within the extracellular cavity of β_2_AR. These results imply that in the late intermediates of the analyzed trajectory (#18-20), the overall disappearing receptor densities are primarily due to the flexible disposition between receptor and G protein, without the receptor undergoing major conformational changes within this timeframe.

In early cryo-EM intermediates (#1-16), the α5 helix is fully engaged and Gα forms interfaces with ICL2, TM5, and TM6 of the receptor. Phe139^34.51^ on the ICL2 of β_2_AR makes contacts with Phe362, Arg366, and Ile369 on the α5 helix, and with His41 on the αN-β1 hinge loop (Fig. 5a). The immediately adjacent Pro138^34.50^ on ICL2 produces an additional α5 contact and participates in coordinating Phe139^34.51^. On the other hand, TM5 makes extensive contacts with Gα’s C-terminus, α5 helix, α4-β6 loop, and α4 helix, while TM6 primarily contacts the C-terminal residues of Gαs. Remarkably, the majority of these interactions with the receptor are progressively lost as the AHD closes upon the GTP-loaded RHD (#15-20). At the macroscopic level, as evident when all models are aligned by the receptor structure, the G protein heterotrimer assumes small but increasing counterclockwise rotations (as viewed from the cytoplasm) across the receptor axis over the trajectory intermediates, suggesting that the pathway of G protein disengagement from the receptor is directional (Supplemental Fig. 8). This in-plane rotation may be important to destabilize interactions with TM5, which appears to extend its cytoplasmic helicity only upon establishing interactions with the RHD of Gα. Disengagement of G protein from β_2_AR would be a logical next step following changes at the interface of Gαs and β_2_AR that occur in later intermediates (#18-20), particularly given the dramatic restructuring of the Gα α5 helix and C-terminus, which form the central point of contact with the receptor.

**Figure 5.**
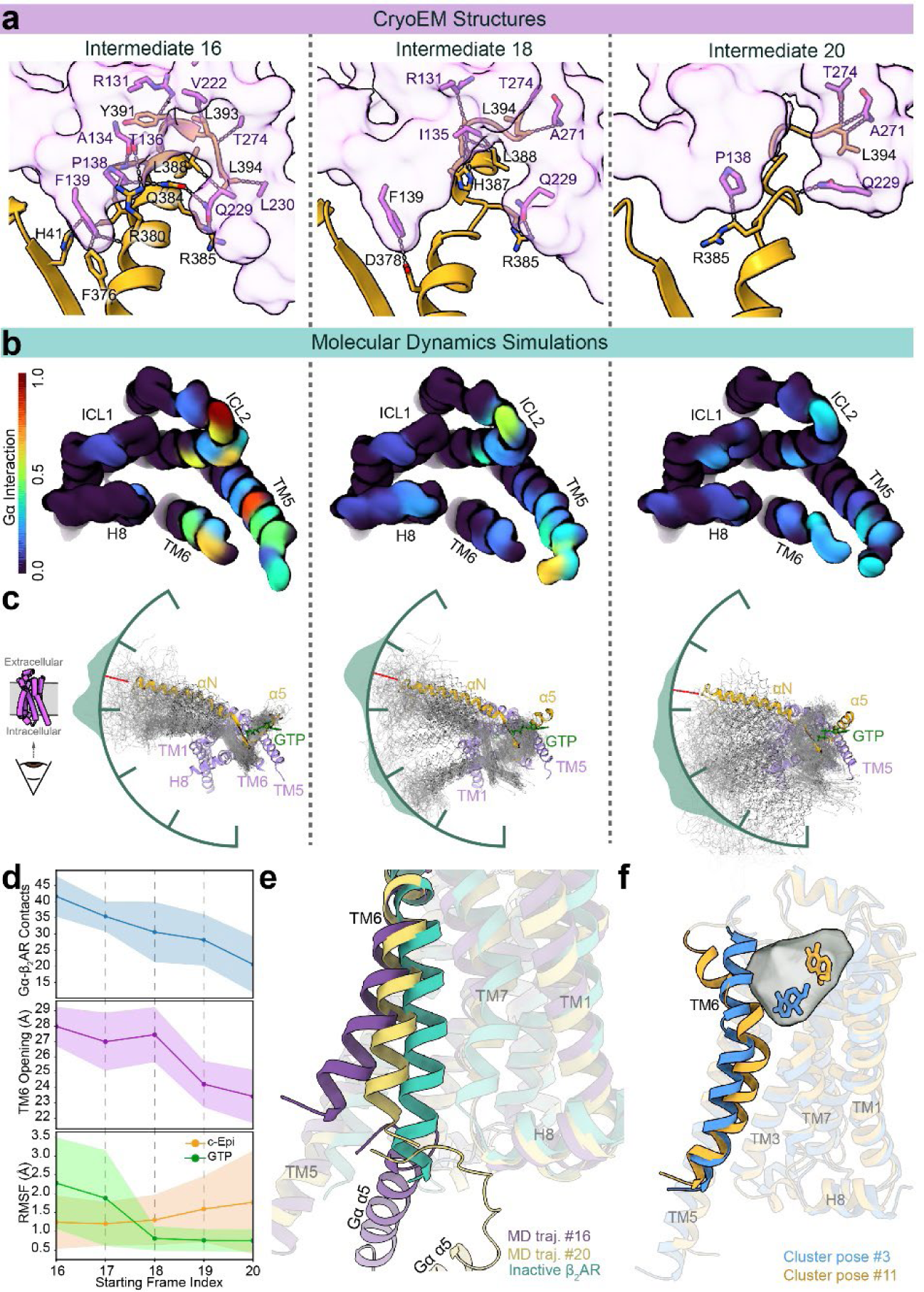
Destabilization of the β_2_AR-Gs Interface. **a**, Interactions between β_2_AR and Gs decrease over the activation trajectory in the cryo-EM structures. **b**, MD simulations starting from the cryo-EM intermediate structures show that the sum of interactions between β_2_ΑR and Gs over the MD trajectories decrease with starting structures from later cryo-EM frames, particularly at ICL1, ICL2, and TM5 (see also Supplemental Fig. 10a). **c**, The decrease in β_2_AR-Gs interaction coincides with flexibility of the G protein in relation to the receptor. MD models were aligned to β_2_AR, the initial structures for each trajectory are shown in full color with resulting periodic trajectory snapshots overlaid in grey. Encompassing each overlay is the distribution of angles of the Gs in relation to β_2_AR over the MD trajectories. The initial angle is inscribed as red tic. Panels ‘**b**’ and ‘**c**’ are shown as viewed from the cytoplasmic space. **d**, Quantification of Gα-β_2_AR contacts (top), TM6 opening (middle), and mobility of GTP and c-Epi (bottom) over the MD trajectories started from sequential frames #16-20. **e**, TM6 is found in a semi-closed conformation in simulations starting from late cryo-EM frames. Shown are the representative structures from MD simulations started from cryo-EM intermediates #16 (purple) and #20 (yellow) superimposed with inactive β_2_AR (PDB:2RH1)^39^. TM6 and the Gα C-terminus and α5 helix are shown in full color. **f**, Two (of 14) representative ligand poses showing the ligand dynamics captured in the MD trajectories. The gray cloud shows the space sampled by the ligand during the simulations (see also Supplemental Fig. 10f). The blue model represents the ligand pose (#3) that is most abundant in trajectories started from earlier intermediate frames, while the orange represents a pose (#11) that develops in MD trajectories started from cryo-EM intermediate #20. The extracellular half of TM7 has been hidden to show the ligand site. TM6 is shown in full color.

### G protein dissociation from the receptor

To further probe the structural transitions in the late steps of β_2_AR-Gs^GTP (Merged)^, we performed molecular dynamics (MD) simulations of intermediate frames #16-20. For this work, we docked the locally refined receptor models into the globally refined density maps to create composite models with more complete receptor information. Triplicate runs for each cryo-EM intermediate structure over 3 μsec simulations revealed a similar, but progressive, sequence of events over the time course of the MD trajectories. GTP was positionally variable over the simulated trajectory arising from cryo-EM intermediate #16-17. Correspondingly, GTP stabilization through enhanced interactions within its binding site increased over the course of the simulations, with the Switch regions migrating towards and progressively increasing their number of contacts with the nucleotide. In the MD trajectories starting from frames in which the ionic interaction between Asp215 on the β2-β3 loop and Arg373 on α5 is still present (intermediate frames #16-17), the interaction is maintained 60-90% of the simulated time. Strikingly, however, this interaction never re-forms in the MD trajectories starting from an already broken bond (intermediate frames #18-20), indicating the propensity of the Asp215-Arg373 interaction to break in the transitional structures (frames #16-17), forming a barrier to complex re-reformation (frames #18-20). This split of the MD data in frames #16-17 vs #18-20 also correlates with an observed destabilization of the interface between G protein and receptor, with a decreasing number of contacts in MD trajectories starting from intermediate frames #17 and #18 (Fig. 5, Supplemental Fig. 10). In particular, the β_2_AR TM5 decreases contacts with the Gαs α5 helix, β6 strand, and the loop between α4 and β6, while the β_2_AR ICL2 loses contacts with Gαs αN, αN-β1 hinge, β1 strand, β2-β3 loop, and β3 strand (Supplemental Fig. 10). This drop in interface contacts is reflected by the enhanced mobility of the G protein relative to the receptor, which again splits sharply between MD trajectories starting from intermediate frames #16-17 versus #18-20. Notably, a counterclockwise rotation of the G protein relative to the receptor when viewed from the cytoplasmic side, as also found in our cryo-EM data, was observed as a trend in our MD data (Fig. 5c, Supplemental Fig. 10), supporting the concept of a directional dissociation pathway.

Collectively, the MD simulations show that enhanced contacts with GTP upon tight AHD closure correlates with Gαs α5 helix destabilization and that the structures representing the late frames (#16-20) of the cryo-EM trajectories lead to functional dissociation, an event that becomes increasingly irreversible upon the initial destabilization of receptor-G protein interactions. In one trajectory started from frame #20 we observed complete detachment of the G protein from the receptor. In MD trajectories started from the structures of the latest frames (#19-20), the gradual disengagement of the G protein correlates with the transition of the cytoplasmic half of TM6 towards a close conformation, a trademark of GPCR inactivation that reduces the accessibility of the intracellular receptor cavity to G proteins or Arrestin^39,40^ (Fig. 5d-e). In a lever-like fashion, the inward movement on the intracellular side of TM6 results in an outward movement of its extracellular side (Fig. 5d-e, Supplemental Fig. 10), which correlates with increased mobility of the ligand c-Epi within the ligand binding cavity (Fig. 5f, Supplemental Fig. 10). Characteristically, c-Epi tends to migrate towards the putative entry channel and the extracellular vestibule associated with ligand entry^41^. These results, which reflect the allosteric communication between the extracellular ligand binding pocket and the intracellular G protein binding cavity^40,42^, further reinforce the validity of our findings and suggest that the TM6 of β_2_AR approaches a conformation similar to the inactive-state relatively swiftly upon functional dissociation of the activated G protein.

### Stepwise mechanism of G protein activation by GTP loading

Our time-resolved cryo-EM structures highlight a sequential series of structural transitions underlying G protein activation upon GTP loading (Fig. 6). These conformational changes can be broadly classified into early-, intermediate- and late-phase events. Initial GTP binding is coordinated by interactions with the TCAT and P-loop of Gα, which change their conformation compared to the nucleotide-free G protein. During early phase events, the AHD is in an open conformation away from the RHD allowing initial binding of GTP. In this phase, the bound GTP may gradually increase its number of contacts with the P-loop and TCAT but without any long-range effects on the rest of the RHD. Marking the beginning of intermediate events is the transition of the flexible AHD towards a closed conformation. Unlike the nucleotide-free G protein, the AHD in a closed conformation becomes well-ordered in this state through further interactions with the nucleotide, which essentially bridges the interface between the AHD and RHD. The locking of AHD against the GTP is a watershed event initiating intermediate phase events involving Gα rearrangements. During this phase we observe the helical extension of the α1 helix, presumably due to both the increased coordination of the P-loop by the phosphate tails of GTP and the AHD ordering that connects directly to the α1 via a linker region. We also observe a conformational change in SwII, which comes closer to the γ-phosphate to assume a conformation that correlates with the appearance of Mg^2+^ density in our maps, a result of strong coordination by the β- and γ-phosphates of GTP, Asp223 and the backbone oxygen of Gly226 in SwII, Thr204 in SwI, and α1 Ser54. These events also coincide with the full ordering of the dynamic SwIII towards the nucleotide. The tight stabilization of GTP by the backbone amine of P-loop residues Glu50, Ser51, Gly52; α1 helix residues Lys53, Ser54, and Thr55; and SwI region Arg201 further stabilizes GTP within the nucleotide binding pocket. The stabilized nucleotide also acts to bridge the AHD and RHD through an interaction of Lys293^RHD^ with both the purine ring of GTP and Asp173 of the AHD, while Glu50 and the phosphate tail of GTP interact with Arg201 of the AHD. This full set of GTP interactions marks the beginning of the late-phase events in the activation process.

**Figure 6.**
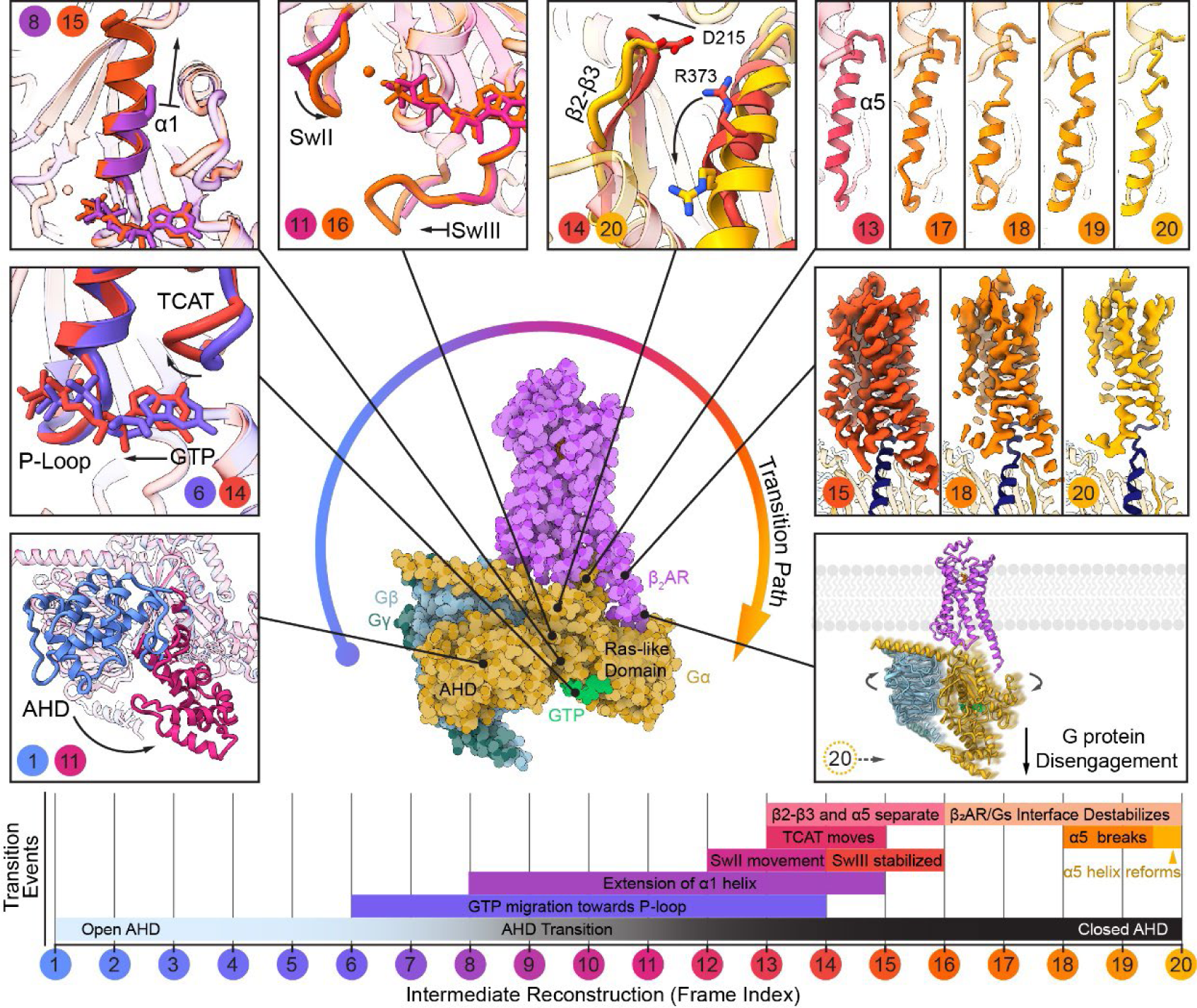
Stepwise activation of G protein following nucleotide exchange initiated by a GPCR. Progression of G protein activation through transitionary events over the course of the 20 cryo-EM structures indicated from 1 to 20 from shades of blue to pink to yellow. Boxed, clockwise from the lower left: Closure of the AHD against the RHD; stabilization of GTP towards the P-loop and corresponding movement of the TCAT motif; extension of the α1 helix; movement of Switch II towards GTP and stabilization of Switch III; distancing of β2-β3 from α5 and breakage of ionic lock; breakage and reformation of the α5 helix into new register beginning closer to the TCAT motif; destabilization of the receptor-G protein complex; disengagement of the G protein from the receptor. Shown in the lower panel is the relative timeline of overlapping events occurring over the cryo-EM trajectory.

Late phase events involve long-range effects of GTP binding with the hallmark of profound structural rearrangements of the α5 helix. These include the unraveling, breaking, and reformation of α5 with a different helical register. Our intermediate frames indicate that Asp215, positioned in the β2-β3 loop, pulls away from Arg373 in α5 due to the interactions of the SwI and SwII loops, flanking β2 and β3, with GTP. The weakening of the Asp215-Arg373 electrostatic interaction appears to allow the partial unraveling of the N-terminal end of α5, likely also due to the strain from the tighter interactions established by the associated TCAT motif with GTP. This allows the reformation of a small helical segment close to the TCAT that appears to grow while helix α5 breaks with extensive unraveling of the C-terminus. The break allows the reformation of α5 with a new register that starts from the helical segment close to the TCAT motif. The destabilization of the “nucleotide-free” conformation of α5 and loss of helical structure at the Gα C-terminus is detrimental to the stability of the interface with the receptor. Late-phase intermediate frames of the cryo-EM trajectory show the deterioration of features in receptor density, the result of flexibility at their interface. The last frame in our reconstruction series reveals no secondary structure at the Gα C-terminus, which has entirely unraveled, giving the impression that the G protein is almost hanging on to the receptor by ‘a thread’. Given the tenuous interactions, we assume that the next step would be the functional dissociation of the G protein, as also fully supported by our MD simulations. Of note, the structure of the Gα C-terminus in the final cryo-EM frame is highly reminiscent of a transition intermediate we previously captured in the cryo-EM structure of the viral GPCR US28 in complex with G11 (PDB:7RKF)^43^, where GDP has not yet been released and the C-terminus of α5 is unraveled proximal to the receptor (Supplemental Figs. 8j-k). This observation supports the notion that the G protein undergoes similar transitions in reverse order to release GDP upon initial association with the receptor.

## Discussion

We developed a time-resolved approach to visualize dynamic events driving G protein activation and receptor disengagement upon GTP binding to a nucleotide-free GPCR-G protein complex. The conformational changes observed in twenty transition cryo-EM structures of pre-steady state β_2_AR-Gs^GTP^ compared to the corresponding analysis of β_2_AR-Gs^EMPTY^ suggest that G protein dissociation upon GTP binding is underlined by ordered structural changes propagating from the nucleotide-binding site and extending to the receptor interface, weakening the interactions between the GPCR and the G protein. Progressive stabilization of the nucleotide between the RHD and AHD correlates with the structural rearrangement of the Gα α5 helix, resulting in destabilization of the receptor interface and the beginning of G protein dissociation. In many ways this process appears to be the inverse to the process of GPCR-G protein association, in which the α5 helix must rearrange outwards to engage the intracellular cavity of the receptor with parallel ejection of GDP. In support of the equivalent conformational pathways involving G protein association and dissociation, a separate MD study examining β_2_AR-Gs protein association found that the process involves an in plane rotation of the G protein against the receptor in the opposite direction to the one we observe here for dissociation. Thus, a corkscrew binding and unbinding pattern appears to underline G protein nucleotide exchange by GPCRs.

The implementation of freeze-trapping at different time points enabled us to monitor the progression of conformational ensembles and confirm our interpretation and ordering of events. For this work, we employed standard equipment to vitrify samples within a few seconds after initiating a “reaction” at 4°C, which was sufficient to monitor and reconstruct a meaningful structural ensemble for the question in hand. However, different kinds of questions or types of complexes may necessitate specialized instrumentation or approaches that can monitor faster kinetics with cryo-EM, including ligand spraying^44^, microfluidic mixing and spraying on grids, as has been demonstrated with ribosomes^45-47^, and also rapid release of caged ligands through laser pulses^48,49^. Likewise, although we found cryoSPARC 3DVA to be suitable for our system, such projects will benefit from a rapidly advancing suite of additional processing tools, such as cryoSPARC 3Dflex^50^, RELION multibody^51^, cryoDRGN VAE^52^, and ManifoldEM^53,54^ to delineate structurally continuous sub-populations among heterogeneous samples.

Beyond providing an enriched mechanistic understanding of G protein activation, we hope that this study provides a powerful demonstration for the orthogonal combination of time-resolved cryo-EM and MD simulations, which can now sample complex structural transitions in realistic computational time scales by starting with cryo-EM structures of pre-steady state conformations. We anticipate that the structural models generated in this and future work will be a valuable resource for developing molecular dynamics simulations using multiple “checkpoint structures” and further combined with machine learning approaches for understanding the structural dynamics of GPCR signaling.

**Supplemental Figure 1.**
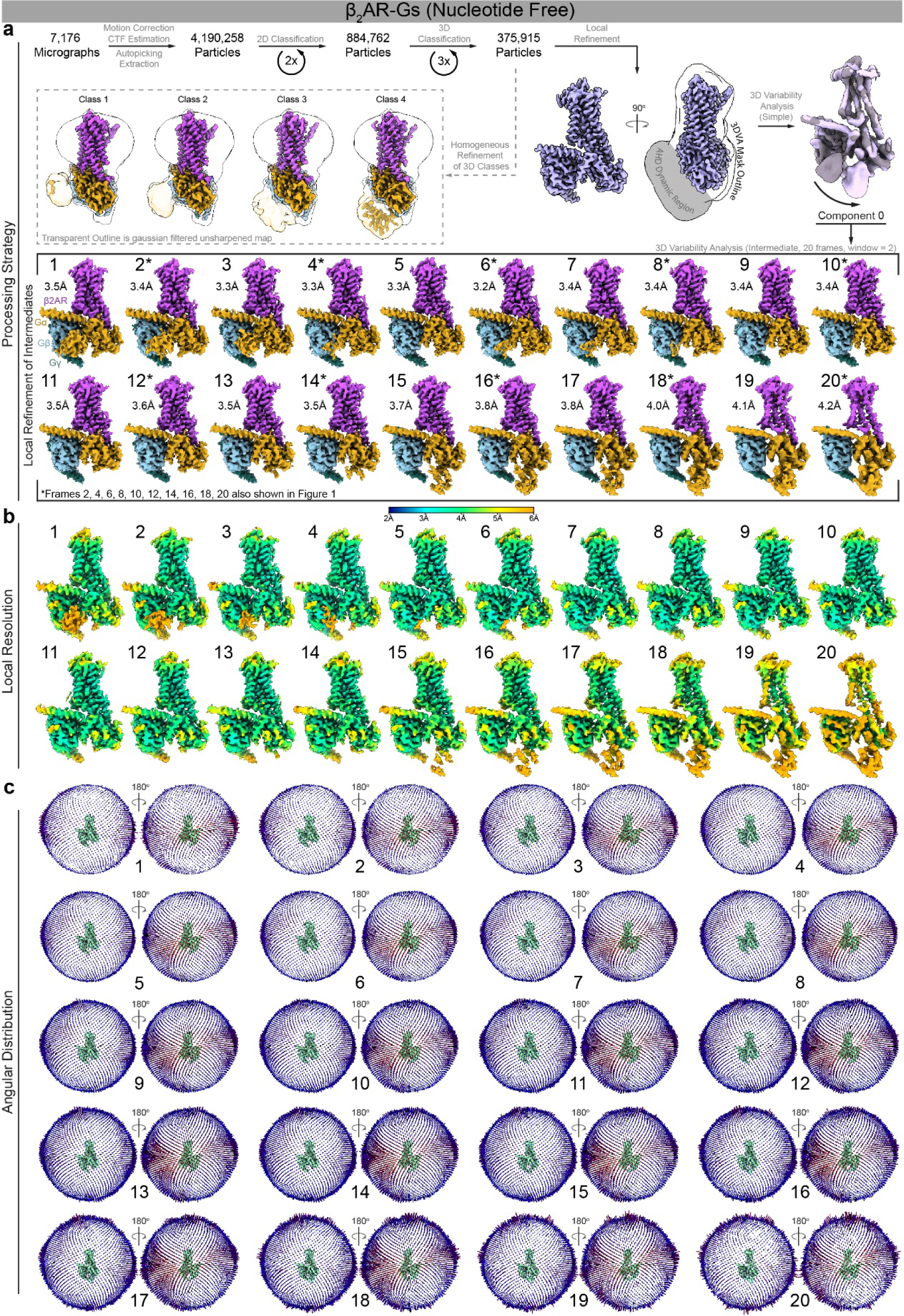
Cryo-EM processing and reconstruction of β_2_AR-Gs^EMPTY^. **a**, Flow chart outlining the cryo-EM processing of β_2_AR-Gs^EMPTY^ complex using cryoSPARC^27,55^. **b**, Local resolution of and, **c**, angular distribution of projections used in final cryo-EM reconstructions.

**Supplemental Figure 2.**
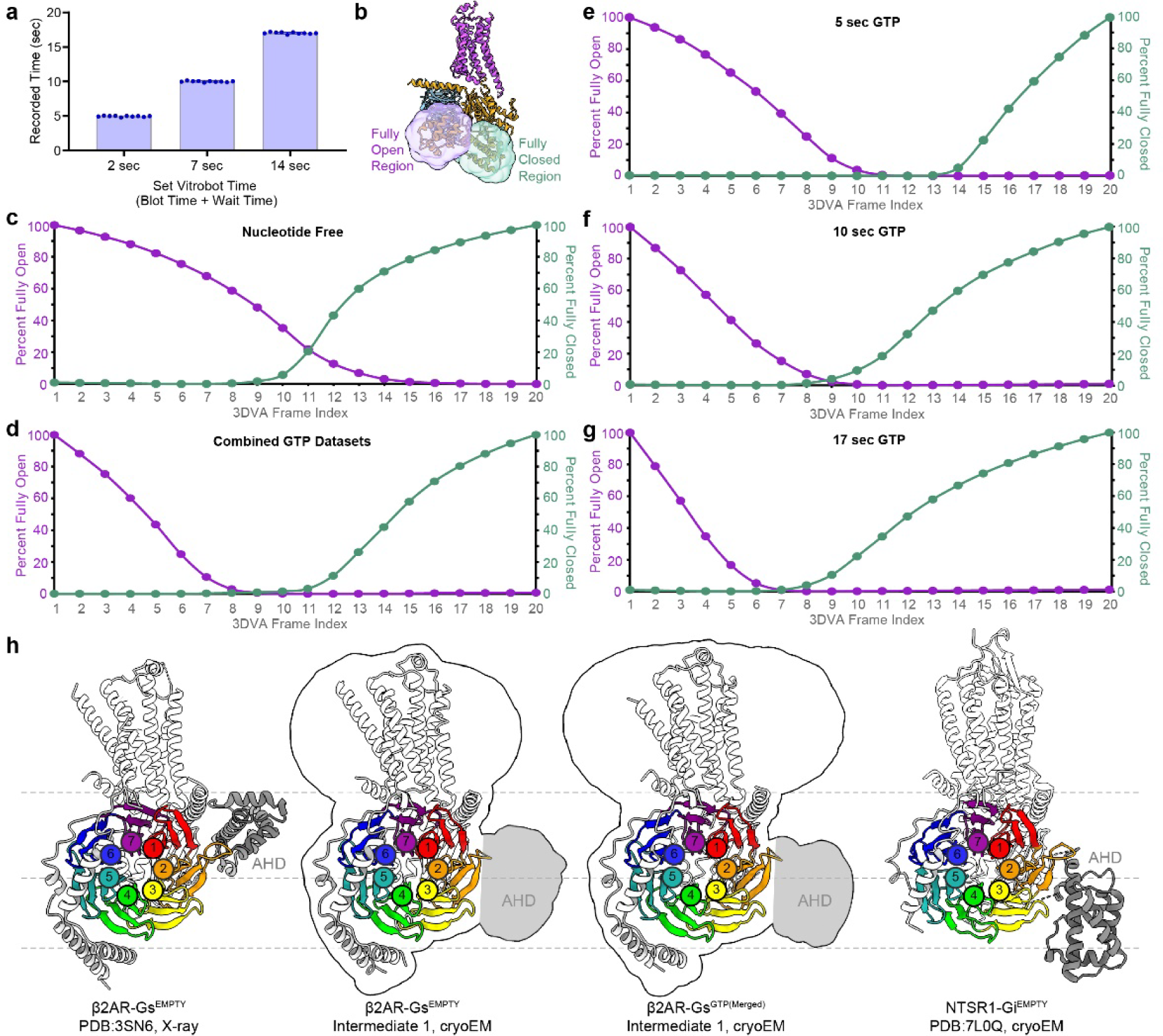
Dynamic residency of Gα AHD in open and closed positions. **a**, Measurement of real time of vitrification using a Vitrobot. Vitrobot timing is the sum of blot time and wait time, 2 sec (4.95 sec ± 0.026 S.E.M., n=10), 7 sec (9.99 sec ± 0.029 S.E.M., n=10), 14 sec (17.02 sec ± 0.040 S.E.M., n=10). **b-g**, To determine the residency of the AHD between open and closed positions in cryo-EM reconstructions, the AHD was docked into frames 1 (maximally open AHD) and 20 (maximally closed AHD) of each 3DVA trajectory, a region of 6Å from the docked structures was used to define ‘fully open’ or ‘fully closed’ respectively, **b**, and the volume of cryo-EM map at a threshold level of 0.05 that was enclosed in the defined regions was determined, **c-g. h**, Location of Gα AHD in relation to Gβ. The crystal structure (PDB:3SN6) locates the Gα AHD (grey) adjacent to Gβ blades 1 (red) and 2 (orange) and interacting with blade 2. In contrast, the location of the cryo-EM density that corresponds to the AHD lies adjacent to Gβ blades 2 and 3 (yellow) in both the nucleotide-free and GTP conditions. The cryo-EM structure of NTSR1-Gi also has an open AHD adjacent to blades 2 and 3, but in a different orientation. Structures have been aligned to Gβ. In the middle panels, the cryo-EM density envelope (gaussian filtered, σ=2) of the unsharpened map is shown with the density corresponding to the location of the AHD shaded in grey.

**Supplemental Figure 3.**
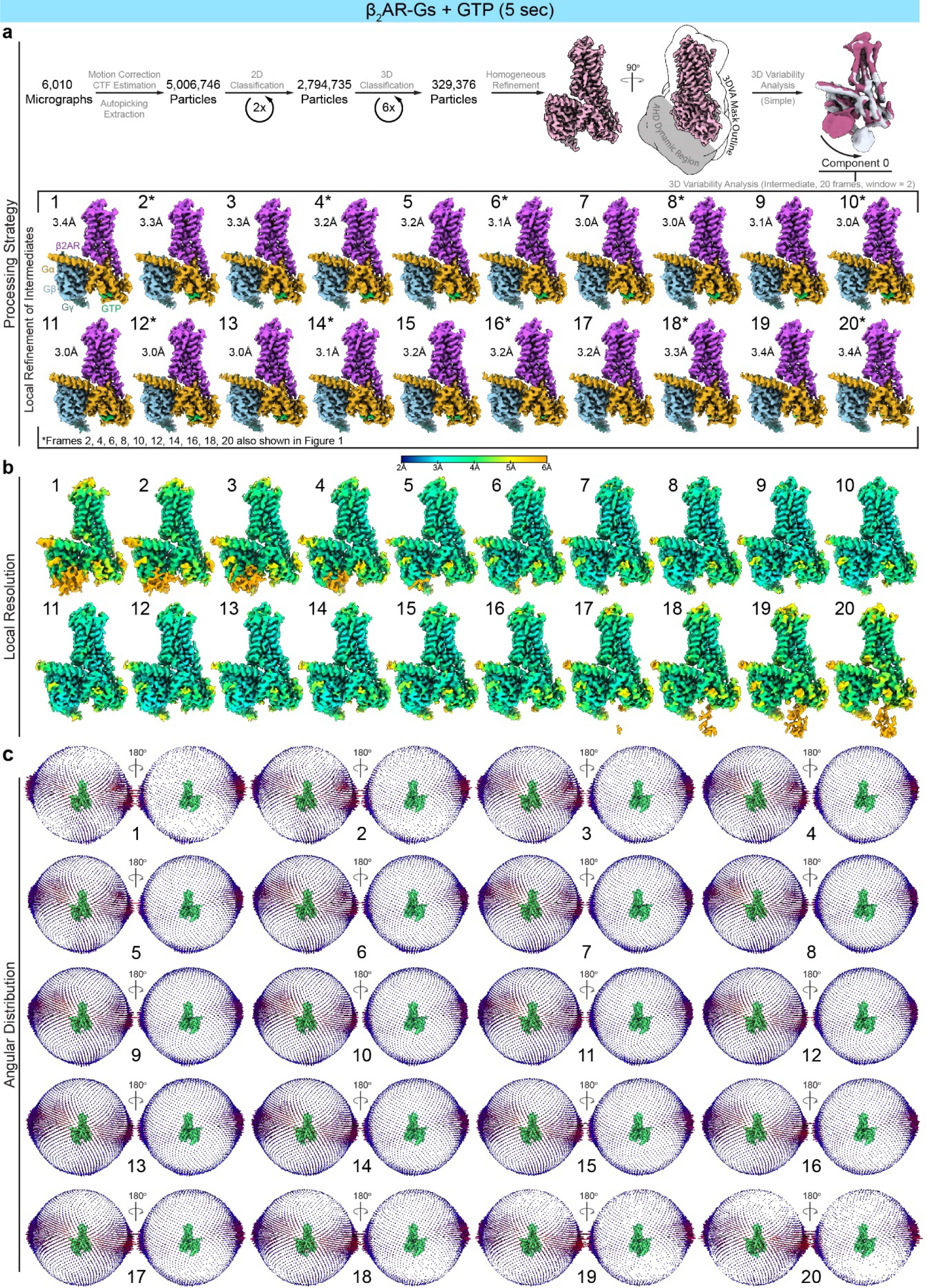
Cryo-EM processing and reconstruction of β_2_AR-Gs^GTP(5sec)^ **a**, Flow chart outlining the cryo-EM processing of β_2_AR-Gs^GTP(5sec)^ complex using cryoSPARC^27,55^. **b**, Local resolution of and, **c**, angular distribution of projections used in final cryo-EM reconstructions.

**Supplemental Figure 4.**
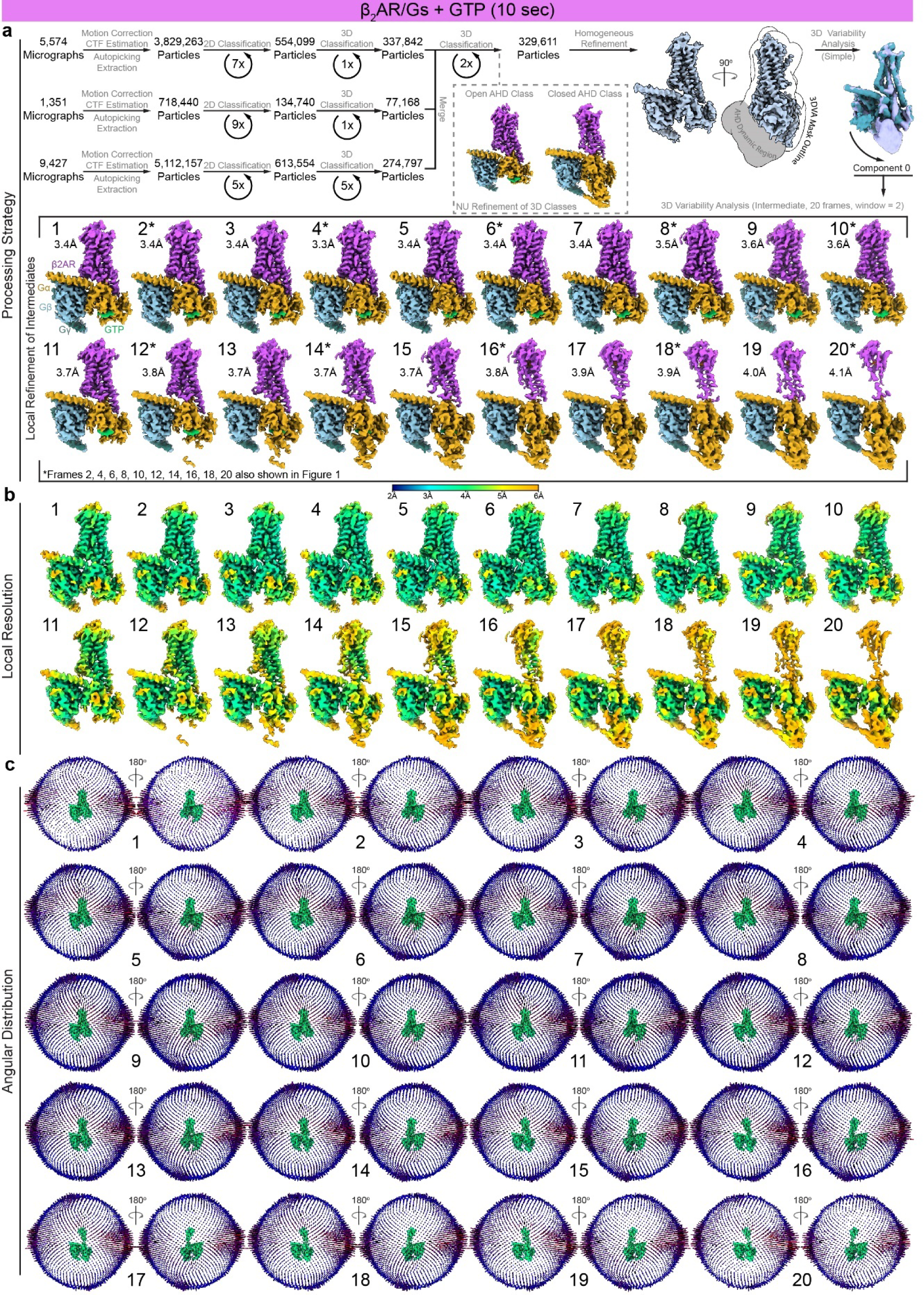
Cryo-EM processing and reconstruction of β_2_AR-Gs^GTP(10sec)^ **a**, Flow chart outlining the cryo-EM processing of β_2_AR-Gs^GTP(10sec)^ complex using cryoSPARC^27,55^. **b**, Local resolution of and, **c**, angular distribution of projections used in final cryo-EM reconstructions.

**Supplemental Figure 5.**
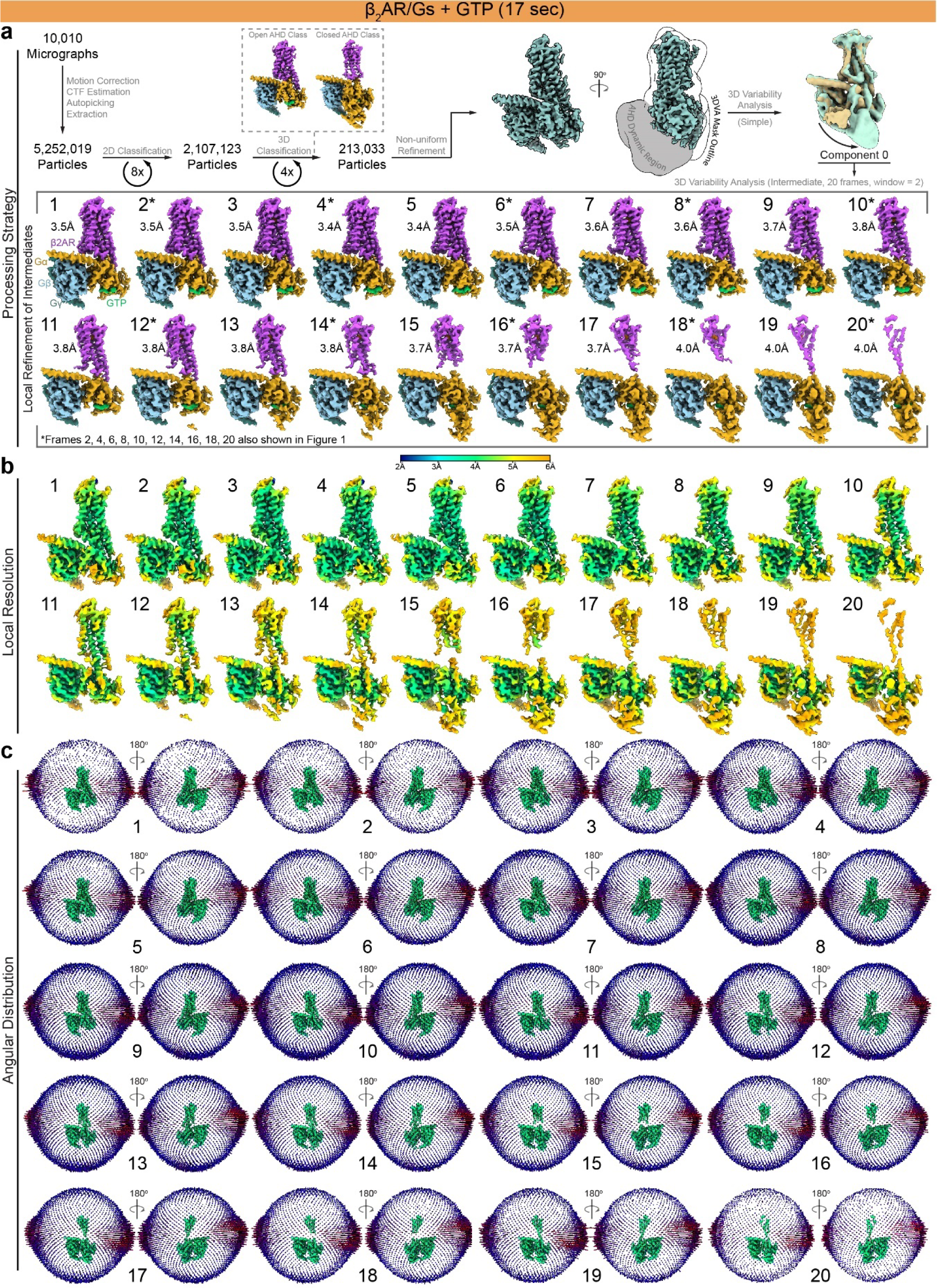
Cryo-EM processing and reconstruction of β_2_AR-Gs^GTP(17sec)^ **a**, Flow chart outlining the cryo-EM processing of β_2_AR-Gs^GTP(17sec)^ complex using cryoSPARC^27,55^. **b**, Local resolution of and, **c**, angular distribution of projections used in final cryo-EM reconstructions.

**Supplemental Figure 6.**
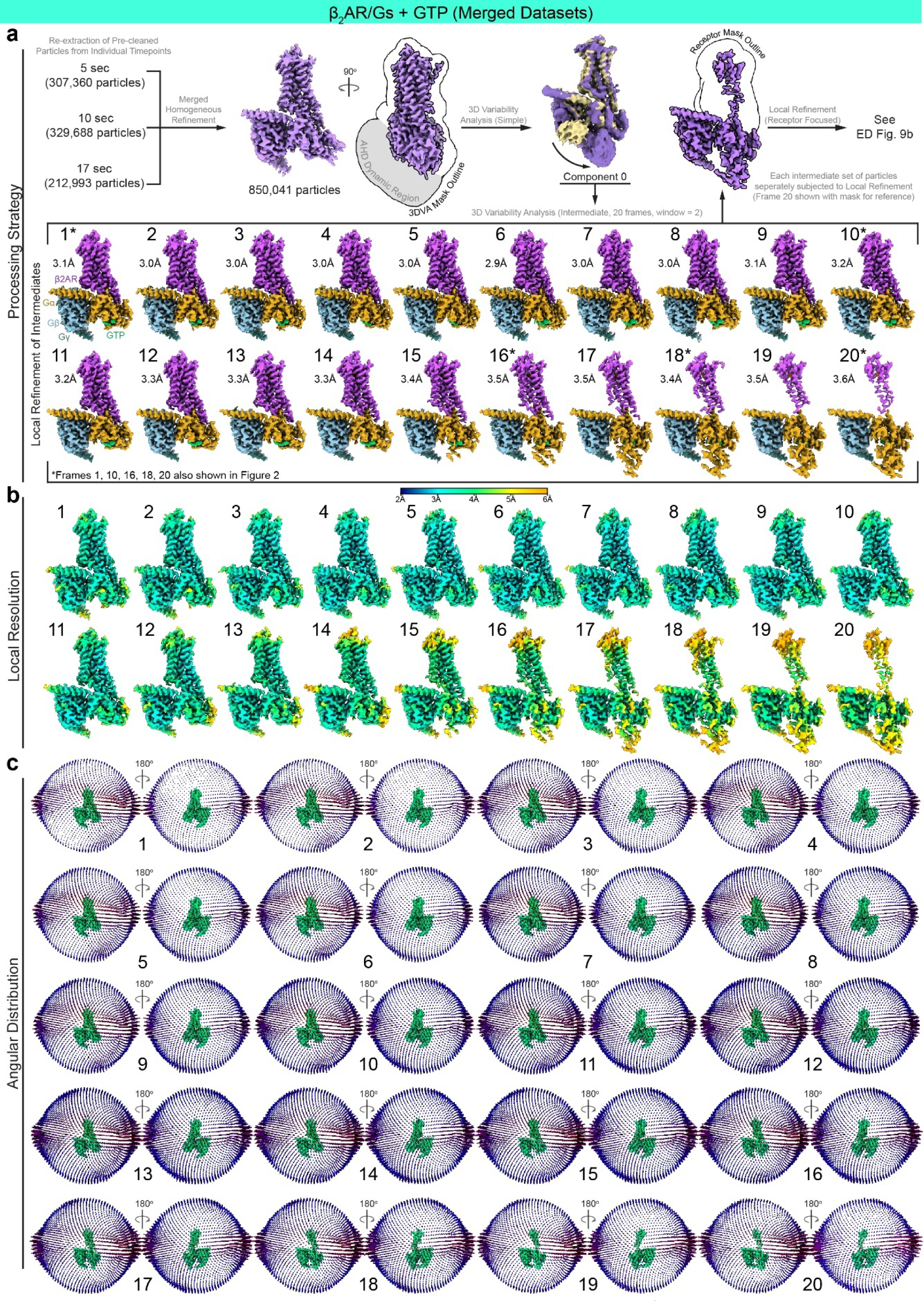
Cryo-EM processing and reconstruction of β_2_AR-Gs^GTP(Merged)^ **a**, Flow chart outlining the cryo-EM processing of β_2_AR-Gs^GTP(Merged)^ complex using cryoSPARC^27,55^. **b**, Local resolution of and, **c**, angular distribution of projections used in final cryo-EM reconstructions.

**Supplemental Figure 7.**
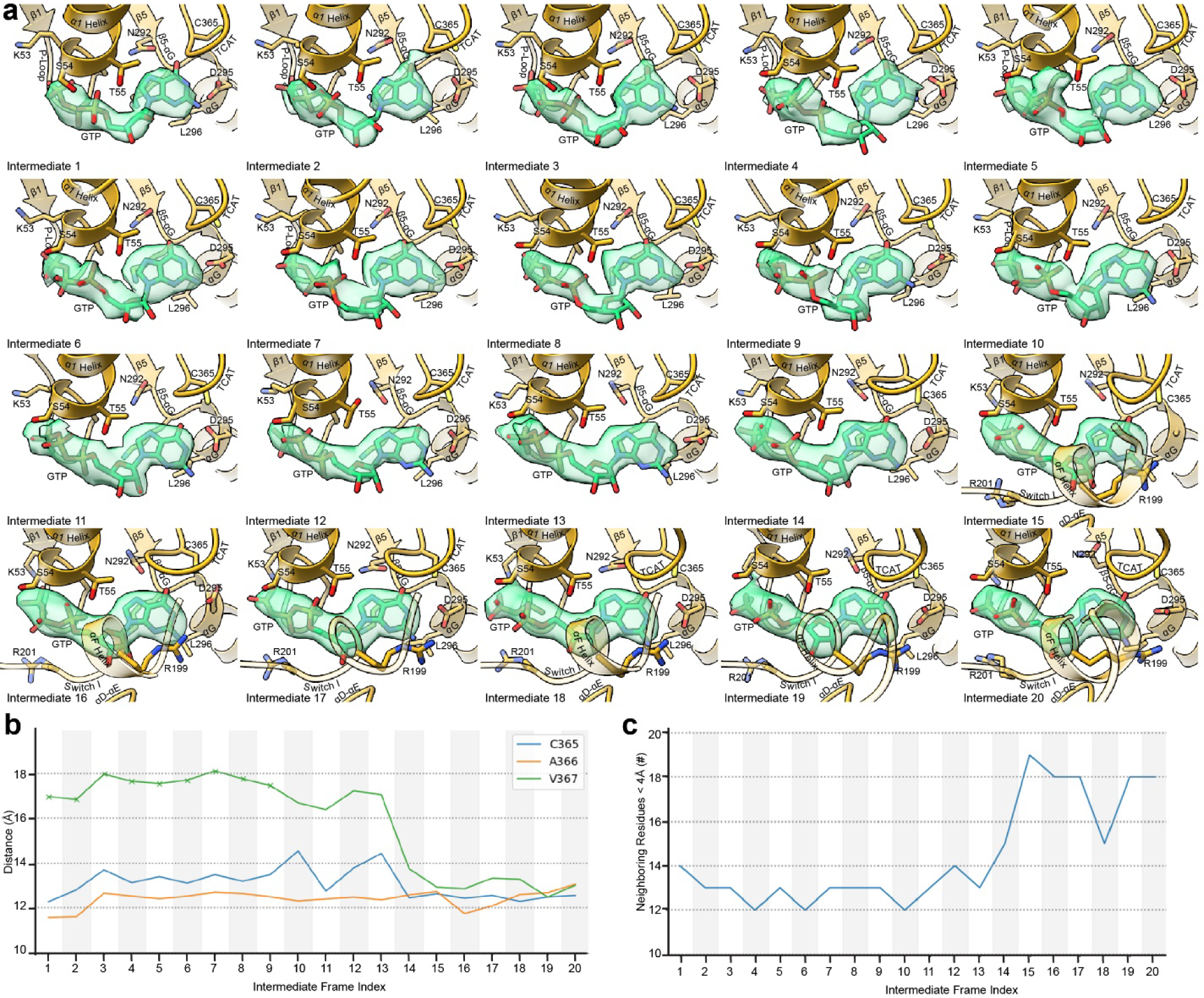
Increased stabilization of GTP in the nucleotide pocket through P-loop interactions with the phosphate tails. **a**, Models showing nucleotide binding site on Gαs across β_2_AR-Gs^GTP(Merged)^ 3DVA trajectory. Cryo-EM density for modeled GTP is shown in translucent green. **b**, For each intermediate frame of the β_2_AR-Gs^GTP(Merged)^ 3DVA trajectory, the traces show selected residue-residue distances between GTP and interaction partners from the TCAT motifs. The marker ‘x’ has been used to indicate residues modelled without their sidechain. **c**, Total Gαs residues in contact with GTP for each intermediate frame of the β_2_AR-Gs^GTP(Merged)^ 3DVA trajectory.

**Supplemental Figure 8.**
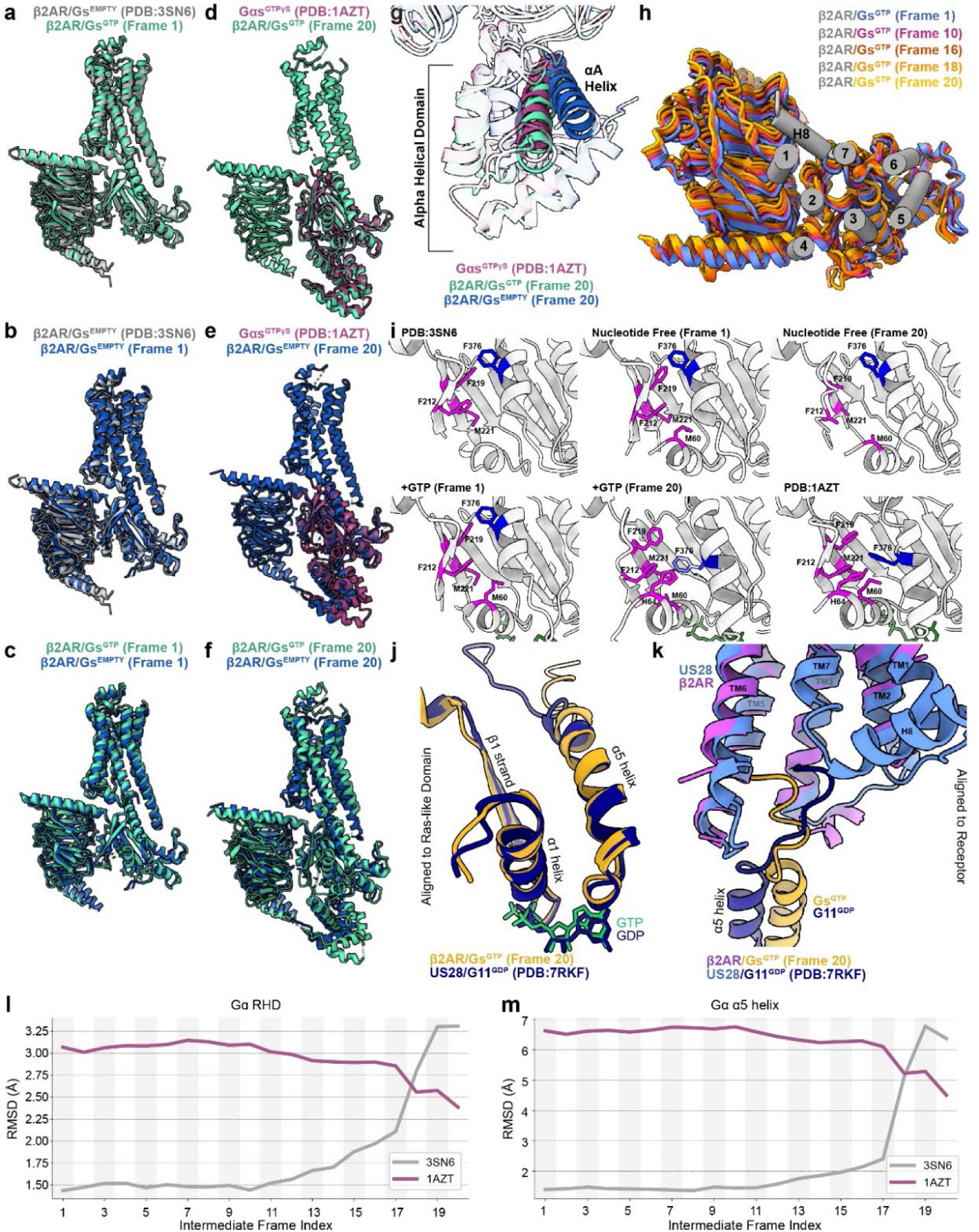
GTP-bound Gαs in the β_2_AR-Gs complex transitions to similar structure as activated Gαs-GTPγS. **a-g**, Structures comparing the overall architecture of the first and last frames of the β_2_AR-Gs^EMPTY^ and β_2_AR-Gs^GTP^ trajectories with ‘checkpoint’ crystal structures of nucleotide free β_2_AR-Gs complex PDB:3SN6 and activated Gαs-GTPγS. Models are aligned to the RHD. **h**, Rotation of Gs in relation to receptor (aligned) over structures of β_2_AR-Gs^GTP^ cryo-EM structural transition frames. **i**, Placement of α5 Phe in relation to hydrophobic pocket on RHD β-sheets. Rendering style inspired by Jang *et. al*.^18^. **j-k**, The transition state of US28-G11^GDP^ captured in the process of nucleotide release is similar to that of β_2_AR-Gs^GTP^ (frame 20). **l-m,** Trace of the root-mean-square-deviation (RMSD) over the 20 β_2_AR-Gs^GTP^ structural transition frames. Structures have been aligned to the rigid elements of the Gαs-RHD, and the RMSD has been computed both for the Cα atoms of the whole Gαs-RHD (**l**) and just of the α5 helix (**m**). The traces show that for both the Gαs-RHD as a whole as well as the α5 helix, the early frames are structurally closer to PDB:3SN6, whereas the last three frames, from 18 onwards, are closer to PDB:1AZT.

**Supplemental Figure 9.**
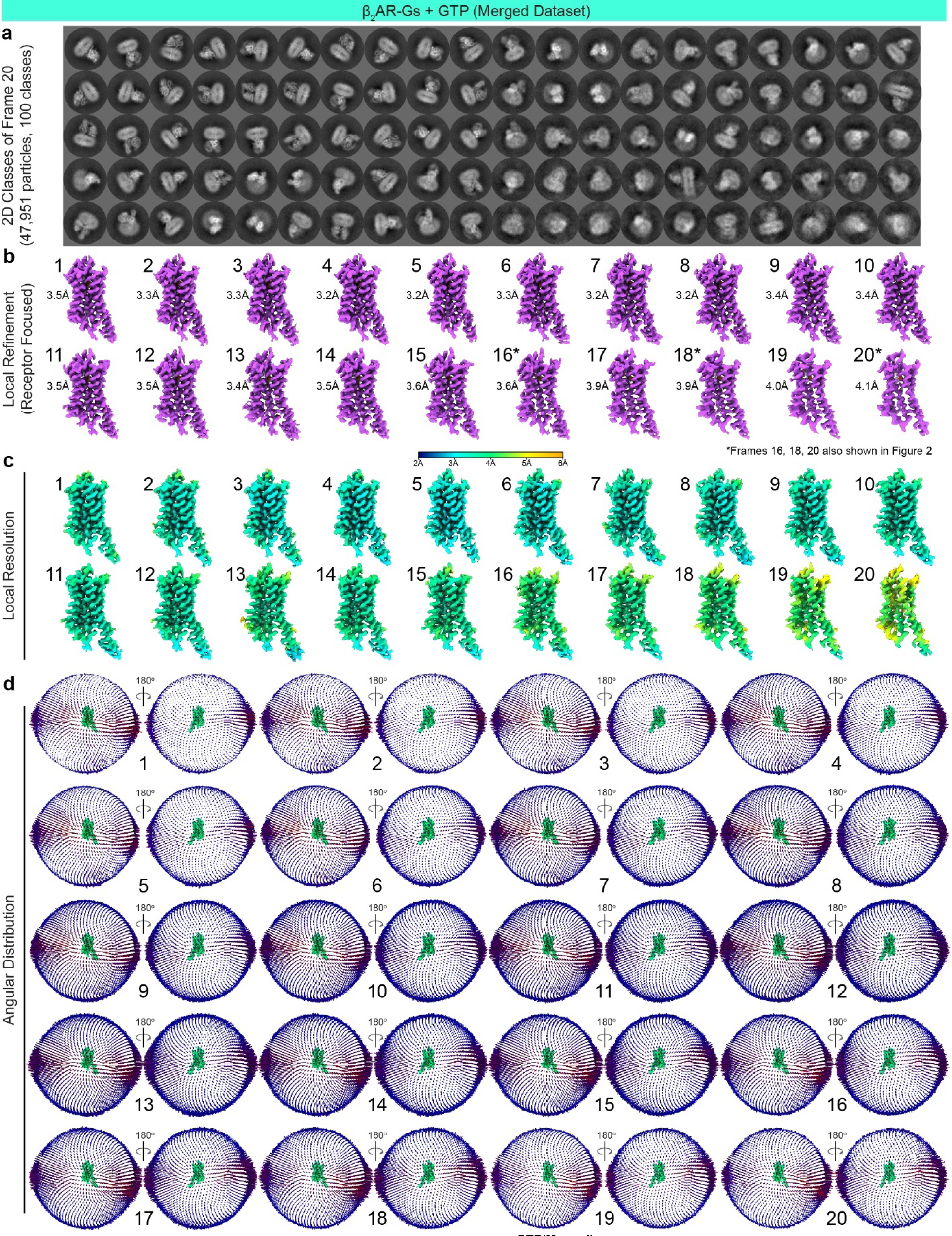
Local refinement of β^2^AR-Gs^GTP(Merged)^ **a**, 2D class averages arising from the 47,951particles contributing to frame 20 of the β2AR-^GsGTP(Merged)^ reconstruction sorted into 100 2D classes. All classes appear to have intact receptor micelle and G protein in complex. **b**, Focused cryo-EM reconstructions of β_2_AR receptor. **c**, Local resolution and, **d**, angular distribution of projections used in final cryo-EM reconstructions.

**Supplemental Figure 10.**
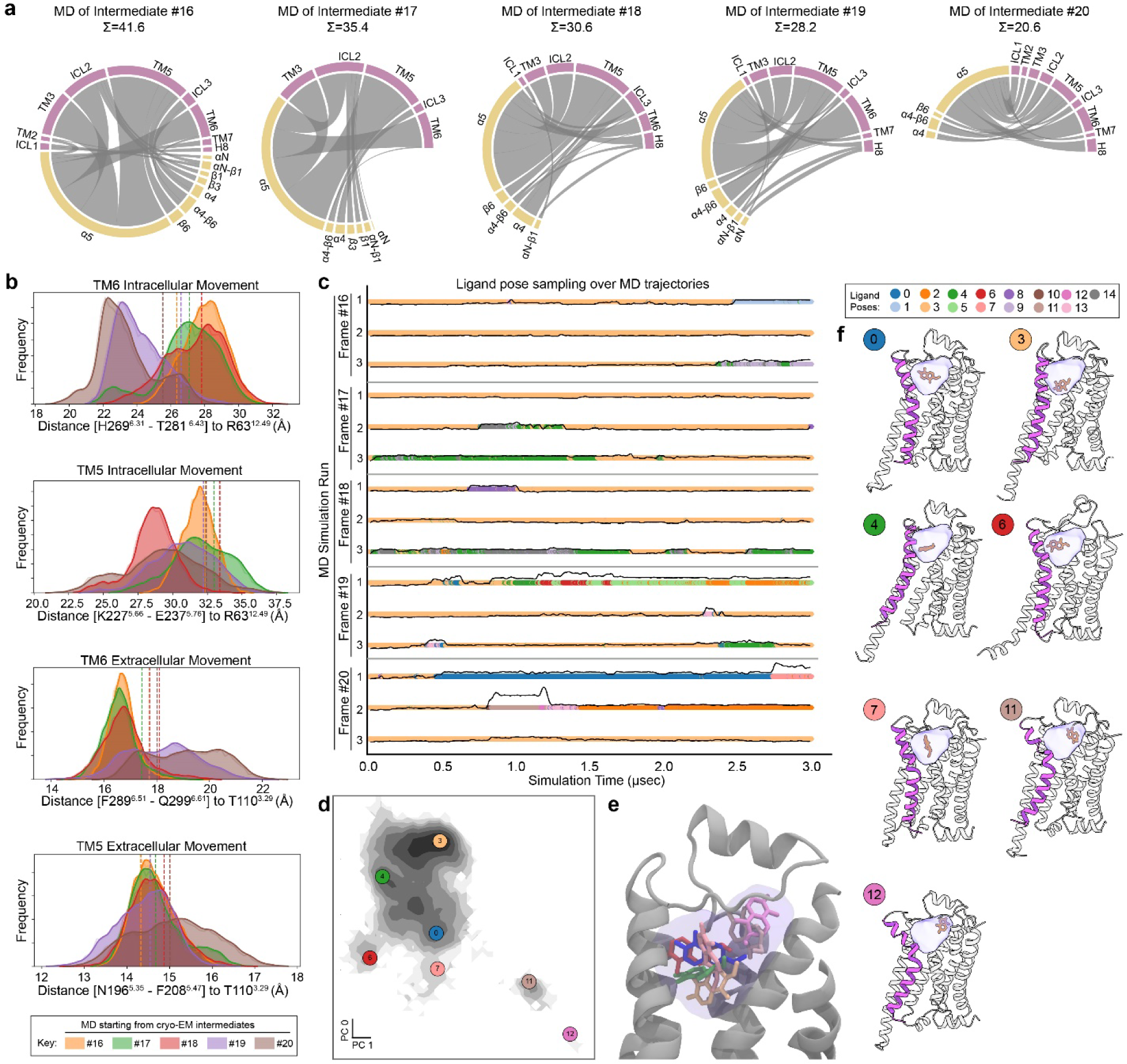
Molecular Dynamics simulations of β_2_AR-Gs^GTP^ intermediate structures. **a**, Weakened interactions of β_2_AR and Gs in simulations seeded by later cryo-EM intermediate structures. Chord diagrams show interactions between receptor regions (purple) with Gα regions (gold) course-grained to fragment segment. Interactions are defined as those less than 4Å and each chord diagram is generated using all the data from three 3μsec MD trajectories for each condition. **b**, Quantification of movement of TM5 and TM6 on the extracellular and intracellular sides of β_2_AR. **c**, Sampling of ligand poses over the MD trajectories shown both as discrete transitions between poses (color-coded time traces, see adjacent ligand pose key above panel ‘f’), as well as in terms of RMSD to the initial pose (solid black line). **d**, Principle component analysis of the sampled ligand poses, with the positions of selected representative poses superimposed as color-coded circled numbers. **e,** Superimposition of selected ligand poses shown in ‘d’. **f**, Representative models of selected ligand pose clusters. TM6 shown in solid purple, c-Epi ligand in orange, and transparent lilac colored cloud represents the area sampled by the ligand across all MD trajectories.

## Methods

### Expression and purification of the β_2_AR for complex formation

β_2_AR was expressed and purified as previously described^1^. Briefly, Sf9 insect cells (unauthenticated and untested for mycoplasma contamination, Expression Systems) were infected with recombinant baculovirus (BestBac Expression Systems) at a density of ∼4.0 × 10^6^ cells per ml. The cells were harvested 55 hr post-infection and lysed by osmotic shock, followed by solubilization of the receptor in *n*-dodecyl-β-D-maltoside (DDM). The soluble fraction was loaded on an M1 anti-FLAG immunoaffinity chromatography as the initial purification step, followed by alprenolol-sepharose chromatography (alprenolol-sepharose resin prepared in-house) to isolate only functional receptors. The eluted receptor was subsequently concentrated on M1 FLAG affinity resin and then washed with ligand-free buffer for 1 hr at room temperature to eliminate the bound orthosteric ligand alprenolol. After elution of the ligand-free receptor with 20 mM HEPES, pH 7.5; 350 mM NaCl; 0.1% DDM; 0.01% cholesteryl hemisuccinate (CHS); 5 mM Ethylenediaminetetraacetic acid (EDTA); and 0.2 mg ml^-1^ FLAG peptide the protein was concentrated in a 100 kDa MWCO Amicon spin concentrator and further purified by size-exclusion chromatography on a Superdex200 Increase 10/300GL (Cytiva) gel filtration column in buffer containing 20 mM HEPES, pH 7.5; 100 mM NaCl; 0.05% DDM; and 0.005% CHS. The monodisperse peak of the receptor was pooled and concentrated to ∼250 μM for further complexing with agonist and G protein heterotrimer.

### Expression and purification of the heterotrimeric G protein G_s_

Heterotrimeric Gs was expressed and purified as previously described^56^. Briefly, *Trichoplusia ni* (*T. ni*) insect cells (unauthenticated and untested for mycoplasma contamination, Expression Systems) were co-infected with two baculoviruses at a density of ∼3.0 × 10^6^ cells per ml, one encoding the human Gαs-short splice variant and the other encoding both the Gβ_1_ and Gγ_2_ subunits, with a histidine tag (6xHis) and HRV 3C protease site inserted at the amino terminus of the β-subunit. Cells were harvested 48 hr post-infection by centrifugation and lysed in a buffer comprised of 10 mM Tris, pH 7.5, 100 μM MgCl_2_, 5 mM β-mercaptoethanol (β-ME), 20 μM GDP and protease inhibitors. The membrane fraction was collected by centrifugation solubilized with a buffer comprised of 20 mM HEPES, pH 7.5; 100 mM sodium chloride; 1% sodium cholate; 0.05% DDM; 5 mM magnesium chloride; 5 mM β-ME; 5 mM imidazole; 20 μM GDP; and protease inhibitors. The soluble fraction was purified using Ni-chelating sepharose chromatography, and the detergent was gradually exchanged from cholate/DDM mixture to 0.1% DDM. The protein was eluted in buffer supplemented with 200 mM imidazole, pooled, and HRV 3C protease was added to cleave the N-terminal 6xHis tag during overnight dialysis in 20 mM HEPES, pH 7.5, 100 mM sodium chloride, 0.1% DDM, 1 mM magnesium chloride, 5 mM β-ME and 20 μM GDP. The cleaved 6xHis tag, uncleaved fractions, and 3C protease were removed by a reverse Ni-chelated sepharose step. The unbound fraction was dephosphorylated using lambda protein phosphatase (NEB), calf intestinal phosphatase (NEB), and Antarctic phosphatase (NEB) in the presence of 1 mM manganese chloride at 4°C for 1 h. Fully geranylgeranylated G_s_ heterotrimer was isolated using a MonoQ 10/100 GL column (GE Healthcare). After binding the protein to the column in buffer A [20 mM HEPES (pH 7.5), 50 mM sodium chloride, 1 mM MgCl_2_, 0.05% DDM, 100 μM TCEP, and 20 μM GDP], the column was washed with buffer A and the G protein heterotrimer was eluted with a linear gradient of 0–50% buffer B (buffer A with 1 M sodium chloride). The main peak containing isoprenylated G protein heterotrimer was collected and the protein was dialyzed into 20 mM HEPES, pH 7.5, 100 mM sodium chloride, 0.02% DDM, 100 μM TCEP and 20 μM GDP. After concentrating the protein to ∼250 μM, glycerol was added to a final concentration of 20%, and the protein was flash-frozen in liquid nitrogen and stored at −80°C until further use.

### Chemical synthesis of c-Epi

5,6-Dimethoxy-3,4-dihydronaphthalen-1(2H)-one (1.90 g, 9.21 mmol) was dissolved in dry toluene (100 mL) which was degassed with N_2_ for 15 min. To the solution was added AlCl_3_ (6.14 g, 46.1 mmol). The mixture was heated to reflux for 1 h and subsequently cooled on ice. Then, water (30 mL) and 2 M HCl (30 mL) were sequentially added. The precipitate was collected by filtration and washed with water (30 mL). The solid was dried under vacuum to give pure 5,6-dihydroxy-3,4-dihydronaphthalen-1(2H)-one as a pale brown solid (1.15 g, 70%).

Benzyl bromide (2.30 mL, 19.4 mmol) was dissolved in acetone (80 mL) and NaI (2.13 g, 14.2 mmol) was added. After stirring at room temperature for 15 min, K_2_CO_3_ was added (4.46 g, 32.3 mmol), followed by addition of 5,6-dihydroxy-3,4-dihydronaphthalen-1(2H)-one (1.15 g, 6.45 mmol). The mixture was heated to reflux for 2 h. Water (100 mL) was added, the product was extracted with EtOAc (3 × 50 mL) and the combined layers were washed with brine, dried (Na_2_SO_4_) and evaporated. The residue was purified by recrystallization from methanol (40 mL), and residual mother liquor was purified by flash column chromatography (4:1 *n*-hexane/ EtOAc) to give 5,6-bis(benzyloxy)-3,4-dihydronaphthalen-1(2H)-one as a solid (2.09 g, 90%).

5,6-Bis(benzyloxy)-3,4-dihydronaphthalen-1(2H)-one (410 mg, 1.14 mmol) was dissolved in Et_2_O (20 mL) and a solution of bromine (117 µL, 2.29 mmol) in Et_2_O (10 mL) was added to the stirred solution. After 1 h, 50% NaHCO_3_ solution (20 mL) was slowly added, and the product was extracted with further Et_2_O (2 × 20 mL). The combined organic layers were washed with Na_2_S_2_O_3_ (10% aq. solution, 30 mL), brine, dried with Na_2_SO_4_ and concentrated in vacuo to give a mixture of the mono- and α,α-dibromo compounds. The crude product was dissolved in dry THF (10 mL) and cooled on ice. To this solution was dropwise added a solution of triethyl amine (167 µL, 1.20 mmol) and diethyl phosphite (154 µL, 1.20 mmol) in THF (10 mL) over a period of 10 min. After stirring for 16 h, water (20 mL) was added, and the product was extracted with EtOAc (2 × 20 mL). The combined organic layers were washed with brine, dried (Na_2_SO_4_), concentrated, and the residue was purified by flash column chromatography (5:1 *n*-hexane/ EtOAc) to give 5,6-bis(benzyloxy)-2-bromo-3,4-dihydronaphthalen-1(2H)-one as a yellow oil (485 mg, 97%).

5,6-Bis(benzyloxy)-2-bromo-3,4-dihydronaphthalen-1(2H)-one (1.44 g, 3.29 mmol) was dissolved in DMF (50 mL) and cooled on ice. To the stirred solution was added glacial acetic acid (226 µL, 3.95 mmol), then after 5 min, a solution of sodium azide (428 mg, 6.59 mmol) in water (3 mL). After 3 h stirring at 0 °C, water (50 mL) was added, followed by CH_2_Cl_2_ (40 mL), and the product was extracted with further CH_2_Cl_2_ (2 × 30 mL). The combined organic layers were washed with brine, dried (MgSO_4_) and concentrated in vacuo. The oil was then dissolved in Et_2_O (30 mL) and the solution was washed with water (3 × 50 mL), brine, dried (Na_2_SO_4_) and evaporated to crude 2-azido-5,6-bis(benzyloxy)-3,4-dihydronaphthalen-1(2H)-on*e* (1.22 g, 93%), which was could be immediately used for the next reaction step.

2-Azido-5,6-bis(benzyloxy)-3,4-dihydronaphthalen-1(2H)-on*e* (550 mg, 1.38 mmol) was dissolved in 1,2-DCE (20 mL) and LiAlH_4_ (1 M solution in THF, 4.13 mL, 4.13 mmol) was added over a period of 1 h. After 4 h, the reaction was cooled on ice and quenched with water (30 mL). The mixture was further diluted with CH_2_Cl_2_ (50 mL), then filtered to remove solids. The product was further extracted with CH_2_Cl_2_ (3 × 30 mL), and the combined organic layers were washed with brine, dried (Na_2_SO_4_) and concentrated to give 2-amino-5,6-bis(benzyloxy)-1,2,3,4-tetrahydronaphthalen-1-ol as a yellow oil (485 mg, 94%), in approximately 2:3 *cis*/*trans* ratio.

2-Amino-5,6-bis(benzyloxy)-1,2,3,4-tetrahydronaphthalen-1-ol, *cis*/*trans*-mixture (4.00 g, 10.6 mmol, approx. 70% *trans*) was dissolved in anhydrous CH_2_Cl_2_ (100 mL). After addition of *N*,*N*-diisopropylethylamine (3.62 mL, 21.3 mmol), Boc_2_O (4.65 g, 21.3 mmol) was added under a stream of nitrogen and the reaction mixture was thereafter stirred overnight (18 h). It was evaporated and the residue was purified by flash column chromatography (isohexane/acetone 5:1 to 2:1), yielding tert-butyl-((1RS,2RS)-5,6-bis(benzyloxy)-1-hydroxy-1,2,3,4-tetrahydronaphthalen-2-yl) carbamate enriched with the *trans*-isomers (>90%). After recrystallization of the beige-pink solid (toluene/ isohexane 2:1), a white, diastereomerically pure powder was obtained (3.01 g, 60% yield). Small amounts of *trans*-compound can be separated on chiral, preparative HPLC (ChiralPak IC) with acetonitrile as eluent, giving first (*R*,*R*)- and second (*S*,*S*)-enantiomer.

To a solution of tert-butyl-((1RS,2RS)-5,6-bis(benzyloxy)-1-hydroxy-1,2,3,4-tetrahydronaphthalen-2-yl) carbamate (7.00 g, 14.7 mmol) in absolute CH_2_Cl_2_ (150 mL) were added 2-3 drops of dibutyltin dilaurate and subsequently (*R*)-methylbenzyl isocyanate (2.49 mL, 17.7 mmol, *ee* >98%). The clear solution was stirred under nitrogen atmosphere at r.t. for 7 d. It was quenched with 2 M NaOH solution (50 mL, stirring for 30 min), the organic layer was separated and the aqueous layer was extracted again with CH_2_Cl_2_. The pooled, organic fractions were washed with water (2x), dried (MgSO_4_) and evaporated, to give a beige powder in quantitative yield. The crude mixture of tert-butyl-((1R,2R)-5,6-bis(benzyloxy)-1-((((R)-1-phenylethyl)carbamoyl)oxy)-1,2,3,4-tetrahydronaphthalen-2-yl)carbamate was recrystallized from toluene/ isohexane (1:1), allowing the hot and clear solution to cool down slowly over the course of several hours. After complete precipitation, the white powder was filtered under vacuum, washed with isohexane/ toluene (4:1), followed by pure isohexane, yielding a residue consisting of 90% (*R,R,R*)-isomer (5.47 g). After a second recrystallization (toluene/ isohexane 5:1, ∼240 mL of solvent), analytically pure (*R,R,R*)-compound was obtained as a white powder (3.90 g, 85%, yield calc. for single diastereomer).

To a solution of tert-butyl-((1R,2R)-5,6-bis(benzyloxy)-1-((((R)-1-phenylethyl)carbamoyl)oxy) −1,2,3,4-tetrahydronaphthalen-2-yl)carbamate (60 mg, 0.096 mmol) in THF (2 mL) was added 4 M LiAlH_4_ solution in Et_2_O (145 µL, 0.58 mmol, 6 eq.) and the resulting reaction mixture was heated to 85 °C for 1 h. After careful addition of water and extraction with CH_2_Cl_2_ (3x), the combined organic layers were washed with brine, dried over MgSO_4_ and evaporated. The resulting crude solid was purified by flash column chromatography (gradient, CH_2_Cl_2_ to CH_2_Cl_2_/MeOH 9:1) to yield (1R,2R)-5,6-bis(benzyloxy)-2-(methylamino)-1,2,3,4-tetrahydronaphthalen-1-ol as a beige powder (23.1 mg, 62% yield).

To a solution of (1R,2R)-5,6-bis(benzyloxy)-2-(methylamino)-1,2,3,4-tetrahydronaphthalen-1-ol (230 mg, 0.59 mmol) in ethanol (15 mL) was added 10% Pd/C (23.0 mg) and the resulting suspension was stirred under hydrogen atmosphere for 2 h. The mixture was filtered through a syringe filter into 0.3% aqueous TFA (50 mL), and the formed solution was frozen and lyophilized. The crude TFA salt was purified by prep. HPLC (0.1% TFA in water + 3% acetonitrile to 10% acetonitrile in 10 min., 12 mL/min. flowrate, peak eluted at 5.0 min) to give c-Epi ((5R,6R)-6-(methylamino)-5,6,7,8-tetrahydronaphthalene-1,2,5-triol trifluoroacetate) as a white powder (142 mg, 74% yield).

### Preparation of the β_2_AR-G_s_ complex for cryo-EM imaging

The β_2_AR-G_s_ complex was prepared essentially in the same way as described previously^1^ using the agonist c-Epi. Briefly, the receptor was incubated with the agonist c-Epi for 1 hr at room temperature prior to the addition of a 1.2-fold molar excess of purified G protein. The coupling reaction was allowed to proceed at room temperature for 90 min and was followed by the addition of apyrase to generate a stable nucleotide-free complex. After 90 min incubation at room temperature, the complex was diluted in a buffer containing 20 mM HEPES pH 7.5, 100 mM NaCl, 10 μM c-Epi, 1% Lauryl Maltose Neopentyl Glycol (LMNG), and 0.1% CHS to initiate detergent exchange. Afterward, the complex was purified by M1 FLAG affinity chromatography to remove excess G protein and residual DDM. The M1 FLAG resin was first washed with buffer containing 1% LMNG, followed by washes with decreasing LMNG concentrations. After elution of the complex with 20 mM HEPES pH 7.5, 100 mM NaCl, 0.01% LMNG, 0.001% CHS, 5 mM EDTA, 0.2 mg ml^-1^ FLAG peptide, and 10 μM c-Epi, the protein was supplemented with 100 μM TCEP and stored overnight at 4°C. The complex was further purified by size exclusion chromatography on a Superdex200 Increase 10/300GL (Cytiva) in 20 mM HEPES pH 7.5, 100 mM NaCl, 100 μM TCEP, 0.001% LMNG, 0.0001% CHS, and 10 μM c-Epi. With the addition of 2 mM MgCl_2_ in the buffer of complex used for GTP experiments. Monodisperse fractions were concentrated with a 100 kDa MWCO Amicon filter.

### Cryo-EM grid preparation

The nucleotide free β_2_AR/Gαs^EMPTY^ complex sample, 15 mg/ml, supplemented with 0.05%octyl-β-D-glucopyranoside was applied to glow-discharged holey carbon grids (Quantifoil R1.2/1.3). The grids were blotted for 2 sec using an FEI Vitrobot Mark IV (ThermoFisher) at 20 °C and 100% humidity and then plunge frozen in liquid ethane. For the β_2_AR/Gαs^GTP^ complex samples, 16 mg/ml, supplemented with 0.02% octyl-β-D-glucopyranoside was applied to glow-discharged UltrAuFoil holey gold grids (Quantifoil, Au300-R1.2/1.3). GTP was added to the grid at a final concentration of 1mM and the grids were blotted using an FEI Vitrobot Mark IV (ThermoFisher) at 4°C and 100% humidity and then plunge frozen in liquid ethane at set timepoints post addition of GTP, adjusted by changing the total of blot time and wait time on the Vitrobot settings (2, 7, and 14 sec). By measuring in real time, using a stopwatch, the time to freeze between the addition of GTP and ethane immersion we found that Vitrobot settings of 2, 7, and 14 seconds equated to 5, 10, and 17 seconds, respectively, in real-time (Supplemental Fig. 2a).

### Cryo-EM data collection

Cryo-EM imaging of the nucleotide-free β_2_AR-Gs^EMPTY^ complex was performed on a Titan Krios (ThermoFisher) electron microscope equipped with a K2 Summit direct electron detector (Gatan) and post-column energy filter. The microscope was operated at 300 kV accelerating voltage, with a nominal magnification of 130,000 x in counting mode resulting in a magnified pixel size of 1.06Å. Μovies were obtained at an exposure of 1.3 electrons/Å^2^/frame with 40 frames per movie stack and defocus ranging from −1.2 – −2.5 μm. Automatic data acquisition was performed using SerialEM^57^ for all data sets. Cryo-EM imaging of the β_2_AR-Gs^GTP (5sec)^ complex was performed on a Titan Krios (ThermoFisher) electron microscope equipped with a K3 Summit direct electron detector (Gatan). The microscope was operated at 300 kV accelerating voltage, with a nominal magnification of 105,000x in counting mode resulting in a magnified pixel size of 0.8677 Å. Μovies were obtained at a total exposure of 60.48 electrons/ Å^2^ over 63 frames with defocus ranging from −1.0 – −2.0μm. Cryo-EM imaging of β_2_AR-Gs^GTP (10sec)^ complex was performed on four separate grids over three collection sessions. All collections utilized a Titan Krios (ThermoFisher) electron microscope equipped with a K3 Summit direct electron detector (Gatan). The microscope was operated at 300 kV accelerating voltage, with a magnification at camera of 58,679 x in counting mode resulting in a magnified pixel size of 0.8521Å. For the first and second grid, movies were obtained at an exposure rate of 21.13 electrons/ Å^2^/sec with defocus ranging from −0.4 - −2.0μm. The total exposure time was 2.717 sec over 77 frames per movie stack. For an additional collection of first grid, movies were obtained at an exposure rate of 20.95 electrons/ Å^2^/sec with defocus ranging from −0.4 −2.0 μm. The total exposure time was 2.717 sec over 77 frames per movie stack. For the third and fourth grids, movies were obtained at an exposure rate of 30.71 electrons/ Å^2^/sec with defocus ranging from −0.5 - −1.6 μm. The total exposure time was 2.008 sec over 79 frames per movie stack. Cryo-EM imaging of β_2_AR-Gs^GTP (17sec)^ was performed on a Titan Krios (ThermoFisher) electron microscope operated at 300 kV accelerating voltage, and equipped with a K3 Summit direct electron detector (Gatan) and post column energy filter, with a magnification of 105,000 x in counting mode resulting in a magnified pixel size of 0.8677Å. Μovies were obtained at an exposure rate of 32.46 electrons/Å^2^/sec with defocus ranging from −0.4 - −0.9 μm. The total exposure time was 1.999 sec over 79 frames per movie stack.

### Image Processing and 3D Reconstruction

Pre-processing of all datasets was carried out similarly, and all processing was performed using cryoSPARC^55^. Dose-fractionated image stacks were subjected to beam-induced motion correction and dose-weighting using patch motion correction and contrast transfer function parameters for each non-dose weighted micrograph were determined by patch CTF. For the β_2_AR-Gs^EMPTY^ complex, 4,190,258 particles from 7,176 micrographs were extracted using semi-automated particle selection. Subsequently, two rounds of 2D classification and three rounds of 3D classification (coupled *ab initio* and heterogeneous refinement operations) were performed on a binned dataset (pixel size 4.24Å and 2.12Å, respectively). A refined set of 375,915 unbinned particles (1.06 Å/pix) was subjected to homogeneous and local refinement. CryoSPARC’s 3D Variability Analysis (3DVA)^27^ was used to determine conformational heterogeneity in the final data set. The former set of particles was processed by 3DVA with three modes, and a mask encompassing the AHD flexible region. Following 3DVA, the first principal component (PC0) was subjected to Intermediate 3DVA Display processing with a window of 2 which sorted particles into 20 overlapping classes that were subsequently processed by local refinement to mask out the detergent micelle.

For the β_2_AR-Gs^GTP^ ^(5sec)^ complex, 5,006,746 particles from 6,010 micrographs were extracted using semi-automated particle selection. Subsequently, two rounds of 2D classification and six rounds of 3D classification (coupled *ab initio* and heterogeneous refinement operations) were performed on a binned dataset (pixel size 3.471 Å and 1.7354 Å, respectively). A refined set of 329,376 unbinned particles (0.8677Å/pix) was subjected to homogeneous and local refinement. 3DVA was used to determine conformational heterogeneity in the final data set. The former set of particles was processed by 3DVA with three modes, and a mask encompassing the AHD flexible region. Following 3DVA, the first principal component (PC0) was subjected to Intermediate 3DVA Display processing with a window of 2 which sorted particles into 20 overlapping classes that were subsequently processed by local refinement to mask out the detergent micelle. For the β_2_AR-Gs^GTP^ ^(10sec)^ complex, a total of 9,706,318 particles from 16,360 micrographs across the collection of four separate grids were extracted using semi-automated particle selection. Subsequently, the particles from each collection were separately subjected to between 5-7 rounds of 2D classification and 1-5 rounds of 3D classification (coupled *ab initio* and heterogeneous refinement operations) were performed on binned datasets (pixel size 3.408 Å and 1.7042 Å, respectively). The particles were then merged to create a refined set of 689,807 unbinned particles (0.8521Å/pix) were subjected an additional two rounds of 3D classification (*ab initio* coupled with heterogeneous refinement), then homogeneously refined. 3DVA was then used to determine conformational heterogeneity in the final data set. The former set of particles was processed by 3DVA with three modes, and a mask encompassing the AHD flexible region. Following 3DVA, the first principal component (PC0) was subjected to Intermediate 3DVA Display processing with a window of 2 which sorted particles into 20 overlapping classes that were subsequently processed by local refinement to mask out the detergent micelle. For the β_2_AR-Gs^GTP^ ^(17sec)^ complex, 5,252,019 particles from 10,010 micrographs were extracted using semi-automated particle selection. Subsequently, eight rounds of 2D classification and four rounds of 3D classification (coupled *ab initio* and heterogeneous refinement operations) were performed on a binned dataset (pixel size 3.471 Å and 1.735 Å, respectively). A refined set of 213,033 unbinned particles (0.8677Å/pix) was subjected to homogeneous and local refinement. 3DVA was used to determine conformational heterogeneity in the final data set. The set of particles was processed by 3DVA with three modes, and a mask encompassing the AHD flexible region. Following 3DVA, the first principal component (PC0) was subjected to Intermediate 3DVA Display processing with a window of 2 which sorted particles into 20 overlapping classes that were subsequently processed by local refinement to mask out the detergent micelle. The β_2_AR-Gs^GTP^ ^(Merge)^ dataset was comprised of the refined particle sets of the β_2_AR-Gs^GTP^ ^(5sec)^, β_2_AR-Gs^GTP^ ^(10sec)^, and β_2_AR-Gs^GTP^ ^(15sec)^ complex datasets that were re-extracted and the particles from the β_2_AR-Gs^GTP^ ^(10sec)^ dataset Fourier cropped to obtain equivalent pixel size (0.8677 Å/pix). The particles were then homogeneously refined together and processed by 3DVA with three modes, and a mask encompassing the AHD flexible region. Following 3DVA, the first principal component (PC0) was subjected to Intermediate 3DVA Display processing with a window of 2 which sorted particles into 20 overlapping classes that were subsequently processed by local refinement to mask out the detergent micelle. The resulting 20 particle sets were additionally locally refined with a mask encompassing the receptor only. UCSF Chimera^58^, ChimeraX^59^, and Protein Imager^60^ were used for map/model visualization.

### Molecular Modeling

The X-ray crystal structure of β_2_AR-Gαs_s_ (PDB ID: 3SN6)^1^ was used as the initial model for the complex in the open AHD conformation, while a composite of PDB:3SN6 with the Gαs-GTPγS crystal structure (PDB:1AZT) was used to generate an initial model for closed reconstructions. The initial models were placed into respective cryo-EM maps using the Chimera ‘fit-in-map’ function. To improve the modeling, iterative rounds of interactive model adjustment in Coot (version 0.9.8.1 EL)^61^ followed by real-space refinement in Phenix (version 1.20.1-4487)^62^ employing secondary structure restraints in addition to the default restraints were completed. Once confidence in the sidechain placement of β_2_AR was reached for the ligand-binding pocket the GemSpot pipeline^63^ utility of Maestro (Schrödinger) was used to dock c-Epi into the maps, then iterative modeling continued, and the final models generated using Phenix refinement. To generate preliminary models for MD simulations the refined models from the global reconstructions (including receptor and G protein) were amended with the local receptor models generated from local refinement of the receptor alone, and then missing architecture (*e.g.*, AHD) was further built-out into low-resolution density using the unsharpened global map to achieve as close of an approximation to experimental data as possible. These preliminary models were then further prepared for MD simulations as described below.

### Model and Map Analysis

To determine the angle of Gαs AHD opening, models with open and closed AHD were aligned to the Ras domain in ChimeraX^59^. Angle of opening is defined as the angle between the center of mass of the closed AHD (residues 88-202), the RHD (residues 203-394), and the open AHD (residues 88-202). The movement of GTP within the nucleotide binding pocket over the 3DVA intermediates was determined by measuring the average change in distance between the nucleotide purine ring and phosphate atoms of the GTP molecule after structures were aligned to the Gαs RHD. To measure comparative volume of density in open versus the closed conformation the AHD was docked into frames 1 (maximally open AHD) and 20 (maximally closed) of each 3DVA trajectory, then a region of 6Å from the docked AHD structures was used to define ‘fully open’ or ‘fully closed’, respectively. The volume of reconstruction EM density, at threshold volume level 0.05, that was encompassed in the defined regions was calculated using ChimeraX^59^.

### Molecular Dynamics Simulations

The β_2_AR-Gs^GTP^ initial structures were extracted from five intermediate frames (#16-20). In the β_2_AR the C-terminus of TM5 and the N-terminus of TM6 was capped at Arg239 and His269, respectively. In Gs^GTP^, Cys2, Ser2, Ala2 and Leu394, Asn341, Cys68 were capped at the N- and C-termini in Gαs, Gβ, and Gγ subunits, respectively. The CHARMM-GUI builder^64^ was used to model and embed the receptor into a pure 1-palmitoyl-2-oleyl-sn-glycero-3-phosphocholine (POPC) bilayer of approximately 150 x 150 (A°)^2^. The palmitoyl group was added to β_2_AR at C341 and N-palmitoyl was added to Gαs, at Gly2, S-palmitoyl to Gαs at Cys3, and S-geranylgeranyl to Gγ at Cys68. In both β_2_AR and Gs^GTP^, all residues were kept in their standard protonation states suggested by the CHARMM-GUI except Glu122, Asp130, Asp79 in β_2_AR that were protonated^65^. Each system was solvated in a rectangular box of 150 Å side lengths for X and Y and 120 Å for Z with TIP3P water^66^ and a concentration of 0.10 M Na^+^/Cl^-^ ions. The CHARMM36^67^ force field was employed for lipids, proteins, and nucleotide. The CgenFF^68^ generalized force field was implemented to describe the β_2_AR ligand c-Epi. All five β_2_AR-Gs^GTP^ intermediates were energy minimized with the steepest descents algorithm and 1000 kJ mol^−1^ nm^−1^ as the threshold. All systems were equilibrated with harmonic positional restraints applied to lipids and Cα atoms of the protein that were sequentially released in a series of equilibration steps. All non-biased simulations were performed using the GROMACS (2022 simulation package)^69^. The software VMD1.9^70^, NLG^71^, MDsrv^72^, and our own python-based analysis package (mdciao)^73^ were used to visualize and analyze MD simulations. NPT simulations were performed at 310K and 1 bar using the velocity-rescaling ^74^ thermostat and Parrinello-Rahman barostat^75^ with a 2 fs integration time-step. Van der Waals interactions were gradually shifted to zero in the range between 10 to 12 Å. Long-range electrostatic interactions more than the cut-off 12 Å were calculated using PME^76^. Relevant hydrogen bonds lengths were constrained using LINCS algorithm^77^. For all five intermediate frames (#16-20), three independent 3-µs-long NPT production runs were carried out for each system setup, started with different initial velocities.

### Analysis of Molecular Dynamics Trajectories

Analysis of the MD simulation data was carried out using Python Jupyter Notebooks^78^ scripted using the python modules mdciao^73^ and MDtraj^79^ for analysis of molecular simulation data.

## Data Availability

The atomic coordinates of β_2_AR/Gs^EMPTY^ (Frames 1-20), β_2_AR/Gs^GTP(Merged)^ (Frames 1-20), along with the coordinates from corresponding localized maps of β_2_AR from the GTP (Merged) dataset have been deposited in the Protein Data Bank.

Cryo-EM maps of β_2_AR/Gs^EMPTY^ (Frames 1-20), β_2_AR/Gs^GTP(5sec)^ (Frames 1-20), β_2_AR/Gs^GTP(10sec)^ (Frames 1-20), β_2_AR/Gs^GTP(17sec)^ (Frames 1-20), β_2_AR/Gs^GTP(Merged)^ (Frames 1-20), along with the corresponding localized maps of β2AR and localized G protein maps have been deposited in the Electron Microscopy Data Bank.

## Contributions

M.M.P-S. prepared cryo-EM grids, collected and processed cryo-EM data and generated maps, built and refined models, analyzed data, prepared figures, and wrote the manuscript. G.P.H. performed data analysis of cryo-EM models and MD simulations and contributed to figure development. H.B. performed MD simulations and data analysis and contributed to figure development. G.E. prepared cryo-EM grids and collected cryo-EM data. G.E. and D.H optimized conditions to obtain stable complexes for the study. Y.G. prepared complex and cryo-EM grids. D.H. purified and prepared β_2_ΑR-Gs complexes. A.B.S and O.P. collected cryo-EM data. M.C. purified β_2_ΑR and Gs, and prepared β_2_ΑR-Gs complexes. F.H. purified Gs and assisted complex preparation. L.M. synthesized c-Epi. P.G. supervised the synthesis of c-Epi. B.K.K. oversaw protein purification and β_2_AR-Gαs complexation. P.W.H. supervised molecular dynamics studies. G.S. conceived project, oversaw cryo-EM studies, and supervised project. M.M.P.-S. and G.S. wrote the manuscript.

## Acknowledgements

Research reported in this publication was supported by equipment access through the Stanford Cryo-Electron Microscopy Center (cEMc). This work was funded by National Institutes of Health grants R01GM083118 to G.S. and B.K.K. and R01NS028471 to B.K.K, and Deutsche Forschungsgemeinschaft (DFG, German Research Foundation) DFG grants GRK 1910 to P.G. and SFB1423, project number 421152132, subproject C01, Stiftung Charité and the Einstein Center Digital for Future to P.W.H. We gratefully acknowledge the scientific support and HPC resources provided by the Erlangen National High Performance Computing Center (NHR@FAU) of the Friedrich-Alexander-Universität Erlangen-Nürnberg (FAU) under the NHR project p101ae NHR funding is provided by federal and Bavarian state authorities. NHR@FAU hardware is partially funded by the German Research Foundation (DFG) - 440719683.

